# The distribution of particulate organic matter in the heterogeneous soil matrix - balancing between aerobic respiration and denitrification

**DOI:** 10.1101/2024.03.16.585355

**Authors:** Maik Lucas, Lena Rohe, Bernd Apelt, Claus Florian Stange, Hans-Jörg Vogel, Reinhard Well, Steffen Schlüter

## Abstract

Denitrification, a key process in soil nitrogen cycling, occurs predominantly within microbial hotspots, such as those around particulate organic matter (POM), where denitrifiers use nitrate as an alternative electron acceptor. For accurate prediction of dinitrogen (N_2_) and nitrous oxide (N_2_O) emissions from denitrification, a precise quantification of these microscale hotspots is required. The distribution of POM is of crucial importance in this context, as the local oxygen (O_2_) balance is governed not only by its high O_2_ demand but also by the local O_2_ availability.

Employing a unique combination of X-ray CT imaging, microscale O_2_ measurements, and ^15^N labeling, we were able to quantify hotspots of aerobic respiration and denitrification. We analyzed greenhouse gas (GHG) fluxes, soil oxygen supply, and the distribution of POM in intact soil samples from grassland and cropland under different moisture conditions. Our findings reveal that both proximal and distal POM, identified through X-ray CT imaging, contribute to GHG emissions. The distal POM, i.e. POM at distant locations to air-filled pores, emerged as a primary driver of denitrification within structured soils of both land uses. Thus, the higher denitrification rates in the grassland could be attributed to the higher content of distal POM. Conversely, despite possessing compacted areas that could favor denitrification, the cropland had only small amounts of distal POM to stimulate denitrification in it. This underlines the complex interaction between soil structural heterogeneity, organic carbon supply, and microbial hotspot formation and thus contributes to a better understanding of soil-related GHG emissions.

In summary, our study provides a holistic understanding of soil-borne greenhouse gas emissions and emphasizes the need to refine predictive models for soil denitrification and N_2_O emissions by incorporating the microscale distribution of POM.

**Highlights:**

- **N_2_O originates from hotspots at the microscale and thus is largely unpredictable**
- **We combined X-ray CT imaging, microscale O_2_ sensors, and ^15^N labeling to map hotspots**
- **The position of the POM separates enhanced aerobic respiration and denitrification**
- **Distal POM (to air-filled pores) as driver of denitrification (N_2_O+N_2_) in soils**
- **Including POM distribution in models will enhance accuracy for soil GHG predictions**

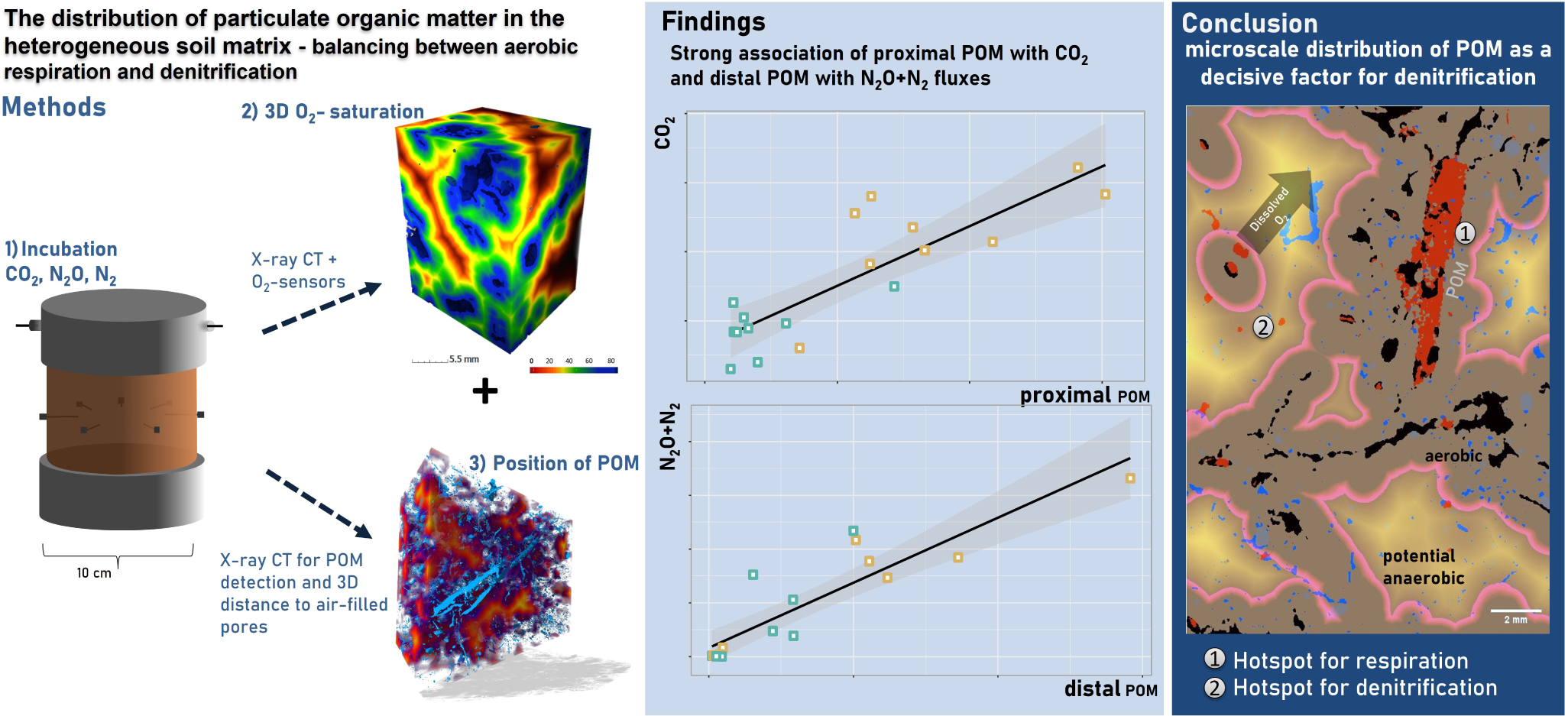

## 1 INTRODUCTION

Nitrous oxide (N_2_O) and dinitrogen (N_2_) emissions from soil significantly impair the nitrogen (N) use efficiency of crops and contribute to global climate change due to the potent greenhouse effect of N_2_O and its role in stratospheric ozone depletion (Ravishankara et al., 2009; Tian et al., 2020). Microbial processes, particularly denitrification, play a pivotal role in N_2_O production, exhibiting complex dependencies on soil structure, moisture, and organic matter content. These dependencies are further complicated by microscale heterogeneity within soils, with completely different microenvironmental conditions at short spatial scales such as oxygen (O_2_) availability or the presence and absence of easily degradable carbon sources. O_2_ availability at the microscale determines the occurrence of either aerobic respiration or anaerobic respiration like denitrification, both releasing carbon dioxide (CO_2_). The latter reduces nitrate (NO_3_^-^) to nitrogen gas (N_2_), with nitrous oxide (N_2_O) as an intermediate (Firestone & Davidson, 1989; Philippot et al., 2007; Robertson, 2023). In contrast, aerobic conditions lead to organic carbon mineralization and concomitant CO_2_ release approximately ten times higher than those under anaerobic conditions due to the higher energy yield of aerobic respiration (Keiluweit et al., 2017; Lacroix et al., 2023). At locations known as hotspots, such as the detritusphere of plant residues, increased C availability leads to intensive microbial activity and thus to increased process rates (Groffman et al., 2009; Kuzyakov & Blagodatskaya, 2015). While, however, the presence of microbial hotspots and their heterogeneous distribution in soil are generally recognized (Kuzyakov & Blagodatskaya, 2015; Lacroix et al., 2023), their quantification remains a challenge.

Yet, a significant advancement in understanding denitrification hotspots, arises from examining the spatial distribution of carbon sources and the O_2_ depletion related to its decomposition. Illustratively, Parkin (1987) found that nearly all denitrification within a soil core could be traced back to a decomposing plant leaf. The persistence and activity of denitrifying enzymes in overall well-aerated soils were therefore attributed to these potential microsites of anaerobiosis around decomposing particulate organic matter (POM, Surey et al., 2020), indicating the nuanced interplay of environmental factors in soils. POM particles, hotspots of oxygen consumption, are distributed non-randomly within the soil matrix (Højberg et al., 1994; Kravchenko & Guber, 2017; Parkin, 1987; Schlüter et al., 2022). This non-random distribution arises for two main reasons: 1) roots predominantly grow into looser areas or directly into macropores (Lucas et al., 2019a, 2022; White & Kirkegaard, 2010) and 2) tillage practices on croplands lead to the vertical mixing of crop residues with soil, which at the same time reduces the population of soil fauna such as anecic earthworms that evenly incorporate POM into the soil matrix (Briones & Schmidt, 2017; Leuther et al., 2023; Wilkinson et al., 2009; Yang & Wander, 1999). Consequently, both processes result in more POM being located near or within macropores.

Oxygen availability at the microscale results from a delicate balance between oxygen consumption through microbial respiration, and supply via diffusion (Keiluweit et al., 2016). In contrast to the distribution of POM, which controls the local O_2_ demand, it is the configuration of the air-filled pore space that determines the diffusion distances of dissolved oxygen through the wet soil matrix (Du et al., 2023; Rohe, Apelt, et al., 2021). This pore space is influenced by processes such as bioturbation (Lucas et al., 2019b; Wilkinson et al., 2009), wetting-drying cycles (Diel et al., 2019; Fomin et al., 2023), or tillage practices (Pires et al., 2017; Schlüter et al., 2018; Wardak et al., 2022). As a result, both O_2_ supply (diffusion distances of dissolved O_2_ through the soil matrix) and local O_2_ demand depend on land use and the associated redox reactions (Lacroix et al., 2023; Lucas, Santiago, et al., 2023; Schlüter et al., 2022).

Historically, the quantification of the anaerobic soil volume fraction and its role in denitrification has been limited by the measurement resolution (Schlüter et al., 2024). For example, estimating anaerobic volume from oxic vs. anoxic incubations provides information on the total anaerobic volume but not the spatial distribution of anoxic microsites. Recent technological advances, such as X-ray computed tomography (CT) imaging, needed to describe diffusion distances to air-filled pores as well as the distribution of POM, and microscale oxygen measurements, to measure local O_2_ concentration, have begun to unveil the complex interplay between soil structure, oxygen dynamics, and microbial hotspots of activity (Kravchenko et al., 2018; Lucas, Santiago, et al., 2023; Rabot et al., 2015; Rohe, Apelt, et al., 2021). Most recently two of these studies revealed that enhanced N_2_O emissions are caused by larger volumes of POM at greater distances to pores, here denoted as distal POM (Lucas, Santiago, et al., 2023; Ortega-Ramírez et al., 2023). In contrast, larger amounts of POM located within or close to pores should result in large CO_2_ emissions from aerobic respiration. However, these authors did not measure complete denitrification (N_2_O+N_2_) and CO_2_ fluxes simultaneously (Lucas, Gil, et al., 2023; Ortega-Ramírez et al., 2023) or incubated homogenized soil (Rohe, Anderson, et al., 2021). Therefore, it is unclear how the position of hotspots created by POM distinguishes between aerobic respiration and anaerobic denitrification in intact soil, which is essential to predict total nitrogen losses and total greenhouse gas (GHG) emissions from soils.

Our study aims to provide a comprehensive understanding of how microscale heterogeneities, specifically the distribution of POM relative to pore structure, influence N_2_, N_2_O and CO_2_ emissions in structured soils. We evaluate results from an incubation experiment with undisturbed soil cores from contrasting environments (cropland vs. grassland) under varying water contents. The combination of X-ray CT imaging with O_2_ microsensors and ^15^N labeling was used for a nuanced exploration of the interplay between soil structure, moisture, and microbial activity, and their collective impact on CO_2_ and N_2_O+N_2_ fluxes. Furthermore, we compare our findings with a previous study using sieved soil without large POM fragments originating from the same sites to understand better the impacts of soil architecture and POM on denitrification processes.

## 2 METHODS

### 2.1 Soil sampling and preparation

In June 2022 undisturbed cores were collected from two sites in Germany representing two land uses and contrasting soils, which were chosen to explore the validity of a universal relationship between microstructure and denitrification for a diverse setting of oxygen demand and supply. The first land use, an arable field, was situated in Rotthalmünster at latitude N48°21’, longitude E13°11’. This field with a maize-wheat crop rotation, undergoes regular plowing to a depth of 30 cm and is subsequently referred to as ‘cropland’. The second land use was an extensively managed grassland (permanent meadow) near Gießen, Germany, which is mown approx. 3 times a year but not fertilized. It is located at latitude N50°32’, longitude E8°41’ and hereafter named ‘grassland’. The soil was a Fluvic Gleysol (IUSS Working Group WRB 2015) with a clay loam texture, containing 4.4% soil organic carbon (SOC) and a pH*_CaCl_* of 6.7, and a bulk density of 1.0 g cm^-3^. The ‘cropland’ soil was a Haplic Luvisol (IUSS Working Group WRB 2015) with a silty loam texture, a pH*_CaCl_* of 7.2 SOC contents of 1.2%, and a bulk density of 1.3 g cm^-3^. Further details can be found in Rohe, Apelt, et al. (2021). For sampling undisturbed cylindrical soil cores, a specialized drill (UGT101 GmbH, Germany) was employed. These samples, encased in rigid 5 mm polyethylene sleeves, consisted of nine undisturbed cores per land use. Each sample measured 10 cm in height and diameter and was extracted from a soil depth of 5-15 cm. The sampling position was chosen randomly in a 10*10 m area without a slope. To reduce the microbial activity to a minimum during storage of the samples, the samples were air-dried until a moisture content of -1000 hPa was reached and stored at 4°C until further processing. At the ‘cropland’, wheat had been harvested just before sampling. Our incubation setup is in accordance with Rohe, Apelt, et al. (2021) In the referenced study, soils were sieved to remove particulate organic matter (POM) to isolate the effects of pore distances on denitrification. Here, we employ the same soil and incubation setup, but with intact field-structured soil, to investigate how POM affects oxygen distribution and thereby influences denitrification hotspots. Note that sieving creates a completely different pore structure (pore sizes and connectivity) and distribution of POM (Lucas, Santiago, et al., 2023) and thus a direct comparison with intact field samples under otherwise identical incubation settings is required to get a complete picture of the influence of microscale heterogeneity on denitrification within the investigated soils. A solution containing ^15^N was created by combining ^15^NKNO_3_ with a 99 atomic percent (^15^N) purity (Cambridge Isotope Laboratories, Inc., Andover, MA, USA) and unlabeled KNO_3_ (Merck, Darmstadt, Germany). The objective was to achieve a nitrogen concentration of 50 mg N kg^-1^ of soil, with the ^15^N-labelled KNO_3_ making up 60 atomic percent of the nitrogen in each level of water saturation treatment. Consequently, appropriate dilution of the stock solution was applied to ensure the same nitrate concentration (per soil mass) and isotopic enrichment in treatments with different soil moisture. In other words, the solution for the different water contents contained exactly the amount of NO_3_ that was needed to retain 50 mg N kg^-1^ of soil in the soil after adjusting the water content. We developed a setup, which assured an even distribution of the ^15^N-KNO_3_ label within the structured cores, as tested with a brilliant blue dye before the experiment. In short, we saturated the samples from below with the labeled NO_3_ solution overnight, then irrigated the samples from above with fresh ^15^N-KNO_3_ solution at a flow corresponding to the saturated conductivity until three times (approx. 1.5h) the pore volume of a cylinder was flushed through the sample. In addition to the uniform distribution of the ^15^N-NO_3_ marker, this ensured that the original soil solution including the dissolved N forms were exchanged. After the flushing, we adjusted the moisture content of the cores by connecting a hanging water column of -8 cm, -40 cm and -160 cm to the bottom of the cores corresponding roughly to the water-holding capacity of 95%, 83%, and 70%. The labeling procedure took place in a 7°C fridge. The water contents at the end of the experiment (corresponding water-filled pore space in parentheses) for -8 cm, -40 cm and -160 cm were 30.2% (71%) ±3.8, 29.1% ±2.8 (65%), 26.5% ±4.9 (54%) for the grasland, and 22.5% ±1.4 (74%), 19.7% ±0.4 (71%), 19.3% ±0.6 (64%) for the cropland, respectively. The two-factorial study design with two land uses, three moisture contents, and three replicates comprised 18 undisturbed soil cores.

### 2.2 Incubation

The columns were closed tightly and equipped with an inlet and outlet in the headspace (Fig. 1). Seven holes were drilled radially in the center of the column to install oxygen microsensors (40×0.8 mm) with *<*140 *µ*m flatbroken sensor tips (NFSG-PSt1, PreSens Precision Sensing GmbH, Regensburg, Germany). The microsensors were attached to a multi-channel oxygen meter (OXY-10 micro, PreSens Precision Sensing GmbH, Regensburg, Germany) to log O_2_ concentration in 30-minute intervals. During the incubation, the columns were placed in separate temperature-controlled (20°C, JULABO GmbH, Seelbach, Germany) and darkened water bath. A constant flow of a technical gas containing 21% O_2_ and 0.5% N_2_ in helium (5 ml min^-1^) was set throughout the 250 h incubation (∽ 11 days) using flow controllers (G040, Brooks® Instrument, Dresden, Germany). This atmosphere with low- N_2_ concentration was used to increase the sensitivity for N_2_ flux measurements (Lewicka-Szczebak et al., 2017). Before incubation started, the headspace was flushed with the technical gas for approx. three hours under three cycles of light vacuum (max. 300 mbar) to reduce the N_2_ concentration in the soil column to approximately that of the technical gas and to ensure comparable initial conditions for the incubation. Our setup allowed the incubation of a block of three parallel samples, i.e. with all three moisture contents simultaneously.

**Figure 1.**
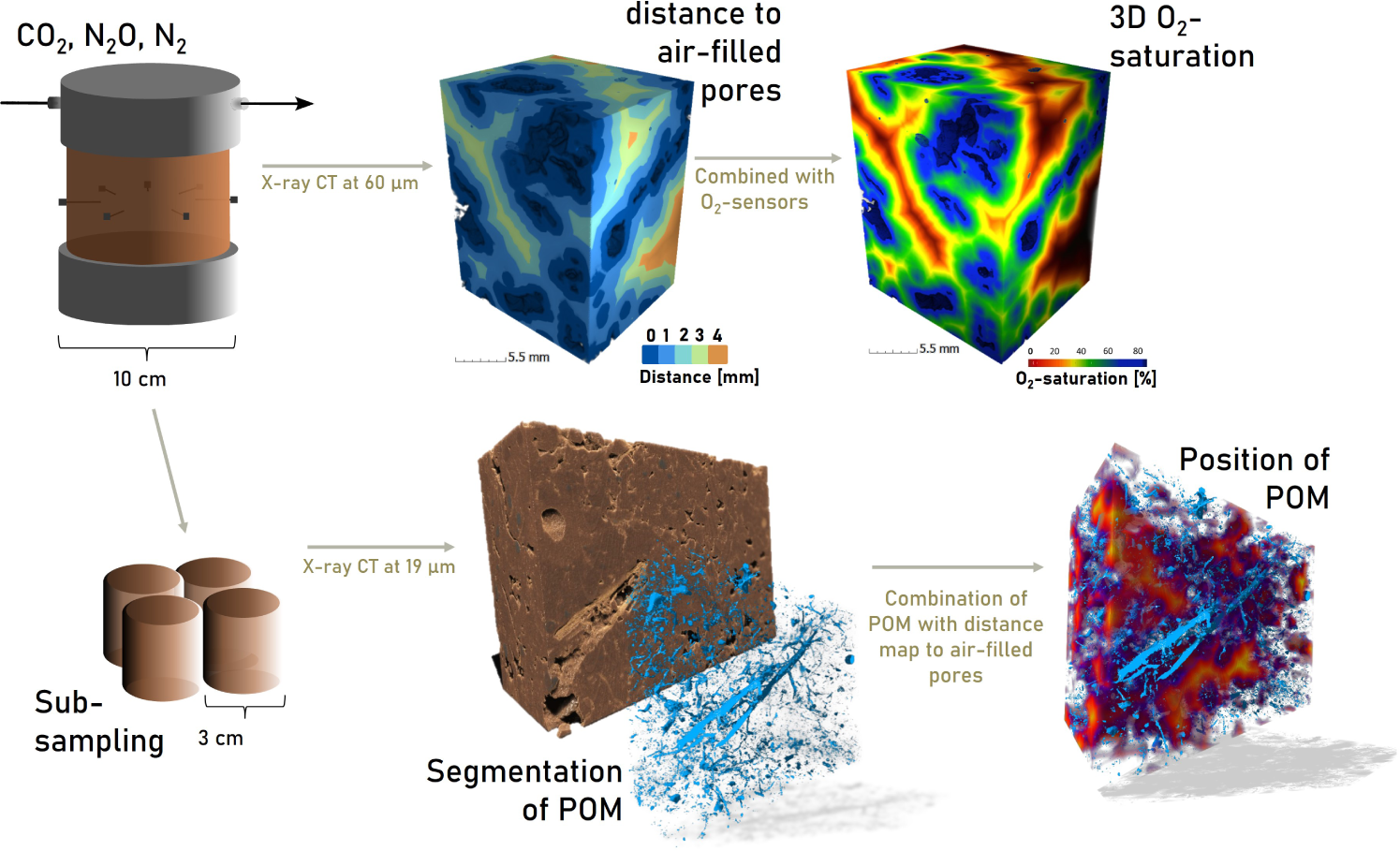
Experimental setup and imaging workflow. The incubation is conducted with 10 cm soil cores under a constant flow of a technical gas with 21% O_2_ and 0.5% N_2_. The entire cores were X-ray CT scanned and analyzed according to the top row, i.e. Euclidean distance maps were derived to examine the exact position of the O_2_-sensor tips. A linear correlation of these distances with the O_2_ measurements allowed us to construct a 3D map of O_2_ saturation. In addition, subsamples of 3 cm size enabled the segmentation of POM at high resolution and to derive its distribution in the heterogeneous soil matrix (bottom row).

### 2.3 Gas analysis

The outlet of the headspace of the columns was directly connected to a gas chromatograph (Shimadzu Nexis GC-2030) with a Barrier Discharge Ionization (BID) detector. This allowed a constant monitoring of CO_2_ and N_2_O (ones per hour). The detection limit was 0.25 ppm N_2_O and 261.9 ppm CO_2_, with a precision of at least 1% and 0.7%, respectively. Note that these N_2_O fluxes derived from GC analysis may include N_2_O from processes other than denitrification and are thus referred to as the total net N_2_O fluxes. To analyze the isotopic composition of N_2_O and N_2_, samples were manually collected into 12 mL Exetainers® (Labco Limited, Ceredigion, UK) at intervals of 1, 2, 5, and 8 days during the incubation period (two samples per time point). To remove residual air, each 12-mL exetainer was purged with helium (Helium 6.0, supplied by Praxair, Düsseldorf, Germany) three times, followed by air evacuation down to 40 mbar. The exetainers were then flushed with headspace gas from incubated columns for 20 minutes, achieving an eightfold exchange of the exetainer’s volume with the gas. Additionally, after the incubation period, a sample of the technical gas was collected to assess the isotopic signature of the carrier gas. These gas samples were analyzed for isotopic signatures with an automated gas preparation and introduction system (GasBench II, Thermo Fisher Scientific, Bremen, Germany; modification based on Lewicka-Szczebak et al., 2013) connected to an isotope ratio mass spectrometer (MAT 253, Thermo Fisher Scientific, Bremen, Germany). This setup was used to measure the mass-to-charge ratios (m/z) of 28 (^14^N^14^N), 29 (^14^N^15^N), and 30 (^15^N^15^N) for N_2_, and thus to simultaneously determine the isotope ratios of ^29^R (^29^N_2_/^28^N_2_) and ^30^R (^30^N_2_/^28^N_2_). We refer to Rohe, Apelt, et al. (2021) for details of measurement and concentrations as well as fractions of different pools (i.e. concentration of N in N_2_O (*fp_N_*_2_*_O_*) or N_2_ (*fp_N_*_2_) and corresponding fraction of N in N_2_O (*Fp_N_2_O_*) originating from ^15^N labeled NO_3_^-^ pool) as well as corresponding ap-values, i.e. the enrichment of ^15^N in the NO_3_^-^pool undergoing denitrification. The product ratio of denitrification [N_2_O / (N_2_O + N_2_)] was calculated for each sample:

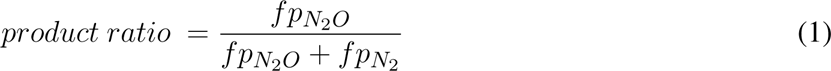

The calculated average product ratio [N_2_O / (N_2_O + N_2_)] of each treatment was also used to calculate the average total denitrification fluxes (N_2_O + N_2_ fluxes) during the incubation in *µg N h^-1^ kg^-1^:*

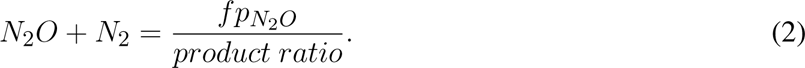

Last, the sum of CO_2_ and N_2_O with a global warming potential of 298 for converting N_2_O to CO_2_-eq. was used to calculate the cumulative GHG production rate as CO_2_-equivalents (CO_2_-eq). We did not consider CH_4_ production, as the GC chromatograms did not indicate any detectable CH_4_ emissions.

### 2.4 X-ray CT scanning and subsampling

After the experiment, the entire 10 cm diameter columns were X-ray CT scanned at 60 *µ*m voxel size (O_2_-microsensors still attached). After this, four intact subsamples with 3 cm diameter and 3 cm height (stiff PVC sleeves with a wall thickness of 2 mm) were taken to be scanned at higher voxel size (19 *µ*m) and thus enabled to segment POM. These subsamples were extracted with a subsampling device (UGT GmbH, Germany) in two depths (2-5 cm and 6-9 cm). X-ray CT was conducted using an X-tek XTH 225 (Nikon Metrology) with an Elmer-Perkin 1620 detector panel (1750 × 2000 pixels). Imaging settings for the large cores were 190 kV, 330 *µ*A, 1.5 mm Cu filter, 2800 projections, two frames per projection, while for the 3 cm cores 130 kV, 150 *µ*A, 0.5 mm Cu filter, 2500 projections, and two frames per projection was used.

### 2.5 Image segmentation and analysis

Image segmentation and analysis were conducted using different plugins in Fiji (Schindelin et al., 2012). In short, the images were segmented into air-filled and water-filled pores, stones, POM and soil matrix using the machine learning plugin LABKIT (Arzt et al., 2022, Fig. S1) along the lines of Lucas et al. (2022). A visual examination of the images indicated no over-segmentation of POM particles. POM particles can only be safely detected if they are thicker than two voxels (38 *µ*m). Nevertheless, according to Schlüter et al. (2022),, these smaller particles appear to be evenly distributed in the soil matrix. Therefore, we assume that analysing the distribution of large POM is sufficient to gain an understanding of the distribution of hotspots. Note that here POM refers to all organic residues visible in the X-ray CT scan, i.e. also relatively intact, but dead root residues cut during sampling. To further elucidate whether the distribution of these intact root segments differs from that of the overall POM, we segmented roots from the entire POM as illustrated in Lucas, Gil, et al. (2023). In short, all POM particles connected to the outer boundary of the image and larger than 10.000 voxels, i.e. approx. 0.06 cm^3^, i.e. these root segments, which still stretch intact through the column, were segmented as intact root segments.

After segmentation, we used the Euclidean Distance Transform (Fig. 1) in Fiji to calculate the potential anaerobic volume, i.e. air distance larger than a threshold similar to Rohe, Apelt, et al. (2021). For this, in the first step unconnected air-filled pores, i.e. those not contributing to a constant delivery of air from the atmosphere, were excluded from the pore image by “connected components labeling” and the “keep largest” method of the plugin MorphoLibJ (Legland et al., 2016, Version 1.4.3) after adding a white slice on top of the image. In this way, all pores connected to the top were kept and all isolated pores were removed. Next, the resulting image was used to calculate the Euclidean distance map, in which each voxel value indicates the 3D distance to the nearest connected and air-filled pore. Last, this 3D map was used to extract the distance threshold for potential anaerobic conditions through sensitivity analyses. This distance divides the soil volume into an aerobic and anaerobic fraction. To derive the optimal distance threshold, we computed linear regressions of the anaerobic volume resulting from varying distance thresholds of 0.06 mm - 5 mm (all soil voxels *>*0.06 mm - 5 mm away from air-filled pores) with the N_2_O+N_2_ fluxes and minimized the *p*-value. We also used the distance maps to calculate the distance of POM to air-filled macropores within the subsamples similar to Ortega-Ramírez et al. (2023) and Lucas, Gil, et al. (2023). The same sensitivity analysis as for the anaerobic soil volume was conducted to estimate the distance threshold. We refer to all POM at higher distances as this threshold as distal POM, while the POM within or close to air-filled pores is denoted here as proximal POM. There was no significant effect of the land use on the distance threshold, so only one regression line was computed for both soils.

### 2.6 Construction of a 3D O_2_-map

The 3D Euclidean distance map of air-filled pores calculated for the large cores was also used to derive the exact positions of the sensor tips relative to air-filled pores within the 10 cm samples. For this, the position of the sensor tips were detected visually and the corresponding values of the distance maps were extracted for each position. After this, we conducted linear regression analysis for the two land uses in which the distance at each of the seven sensor tips per core was correlated with the O_2_ saturation on the last day of the incubation. We used the O_2_ saturation at the end of the incubation to ensure that it aligns best with the structural information derived from X-ray CT scans directly after the incubation. The empirical linear models for O_2_ saturation as a function of air-distance were used to compute the O_2_-saturation of all voxels of the X-ray CT scan, i.e. at all possible distances to air-filled pores (Fig. 1).

### 2.7 ^15^N-NO_3_^-^ and ^15^N-NH_4_^+^ analysis

During subsampling of the larger cores, disturbed soil samples at three different depths (1-3 cm, 5-6 cm, and 8-9 cm) were taken. Fresh soil (10 g) was extracted with 40 ml 1 M K_2_SO_4_solution (corresponds roughly to a ratio of 1:5 solid:liquid in relation to dry matter). After shaking for 1h, the mixture was centrifuged for 30 min at 3000 rpm. The supernatant was analyzed for NH_4_^+^-N and NO_3_^-^-N concentrations and ^15^N-enrichment in NH_4_^+^ and NO3- using the SPINMAS technique (Stange et al., 2007). The measurements were carried out in an automated sample preparatory for inorganic nitrogen (SPIN unit; InProcess, Bremen, Germany) coupled with a mass spectrometer (GAM 400; InProcess, Bremen, Germany). For nitrate analysis, approx. 1 ml V(III)Cl_3_ solution in an acidic medium (HCl) was added to a 5 ml sample at 85°C to reduce the NO_3_^-^ to NO and the m/z values 30 and 31 were measured. To calculate the ^15^N abundance, the m/z value 31 was corrected, due to the interference of ^14^N^17^O (natural abundance of ^17^O 0.037 at%) with ^15^N^16^O. The quantification limit of NO_3_^-^ was 0.005 mmol/L. For analysis of ammonium ca. 1 ml bromate (BrO_3_^-^) in a sodium hydroxide (NaOH) solution was added to a 5 ml sample at 65°C to oxidize NH_4_^+^ to N_2_. To calculate the ^15^N abundance of NH_4_^+^, the m/z values 28, 29, and 30 were measured. The quantification limit of NH_4_^+^ was 0.03 mmol/L

### 2.8 Statistics

For all statistical analysis, the software R 4.3.1 (R Core Team, 2023) in combination with the package agricolae (Mendiburu & Yaseen, 2020) was used. Differences in the data were revealed with a two-factorial ANOVA with the factors of land use (grassland vs. cropland) and moisture content (-160 cm, -40 cm, and -8 cm pressure head). For the fluxes and temperatures, which were logged over time, mean values per core were calculated beforehand. When the interactions between the studied factors were found to be statistically significant (*p <*0.05), Fisher’s Least Significant Difference (LSD) was carried out as posthoc test. The assumptions of normality and homogeneity of variances were assessed using normal probability plots of the residuals and Levene’s tests for equal variances, respectively. To address our research question concerning evaluating soil structural properties as predictors of denitrification, we computed linear regression models of these with the fluxes of N_2_O, N_2_O+N_2_, CO_2_, and GHG.

## 3 RESULTS

### 3.1 Gas fluxes and ^15^N-label

The CO_2_ and total N_2_O emissions over time exhibited considerable variability among the tested soil cores, even among those sharing the same land use type and moisture level (Fig. S2). Notably, there was a distinct land use-dependent influence on CO_2_ emissions, with the grassland demonstrating significantly higher CO_2_ emissions compared to the cropland under drier conditions, particularly at pressure heads of -160 cm (528,3 *µ*g C kg^-1^h^-1^in the grassland compared to 190,0 *µ*g C kg^-1^h^-1^in the cropland) and -40 cm (Fig. 2a, 532,8 *µ*g C kg^-1^h^-1^in the grassland compared to 200,0 *µ*g C kg^-1^h^-1^in the cropland). Conversely, N_2_O emission (Fig. 2b) and combined N_2_O+N_2_ fluxes (Fig. 2c) exhibited an increase with moisture, showing no discernible difference under drier conditions but higher emissions from the grassland under the wettest conditions tested. At the drier pressure heads, grassland samples featured significantly higher product ratios of N_2_O and N_2_ emissions (0.60 at -160 cm and 0.53 at -40 cm pressure head) compared to cropland (0.30 at -160 cm and 0.27 at -40 cm pressure head, Fig. 2d). Yet, these differences diminished, with both land use types showing comparable levels at a pressure head of -8 cm (0.68 for the grassland and 0.63 for the cropland). Thus, at wet conditions, a low amount of N_2_O produced during denitrification was further reduced to N_2_.

**Figure 2.**
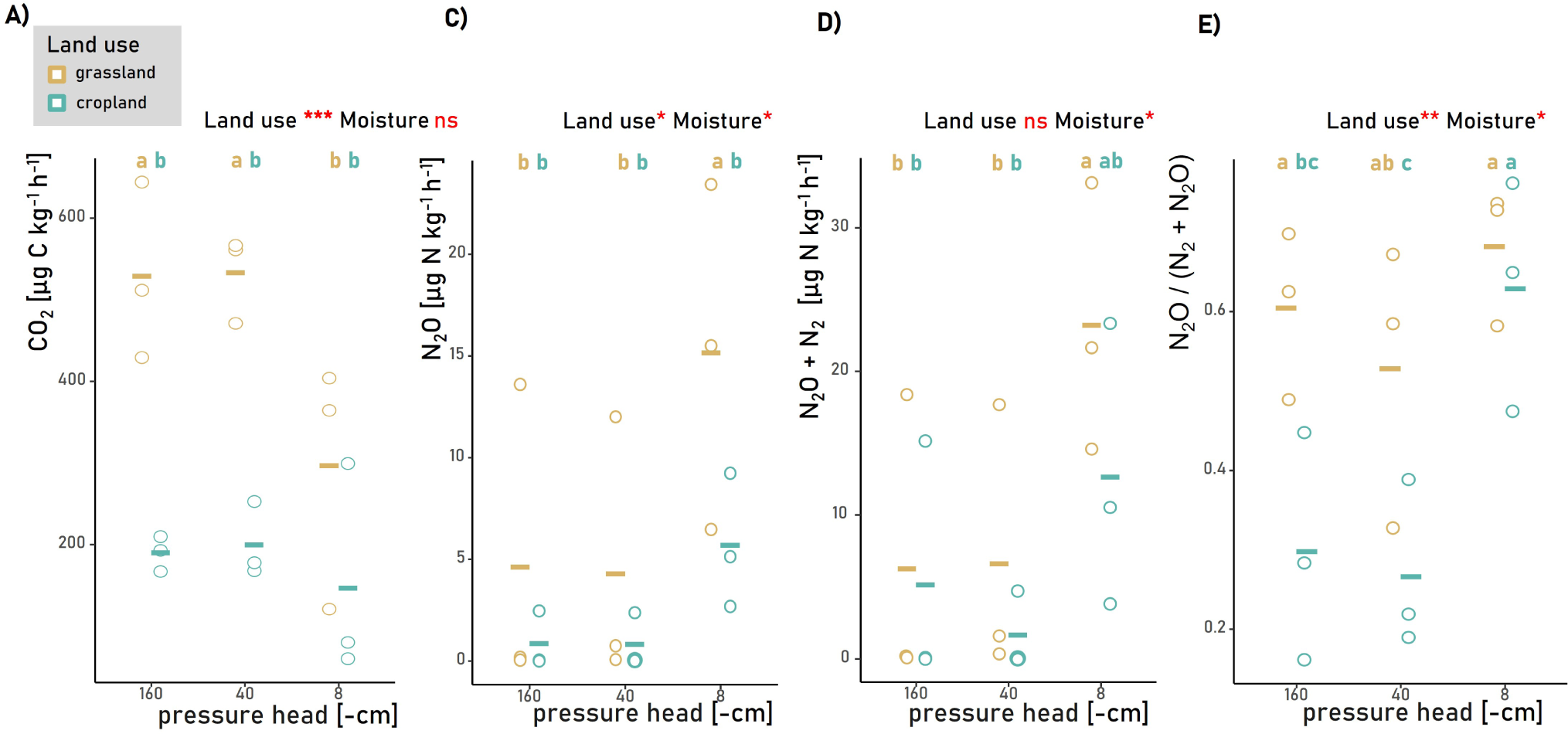
Gas fluxes during the incubation period for the two land uses at three different moisture contents. Shown are the mean fluxes of a core (circles) and the mean of the treatment (crossbar) for A) CO_2_, B) N_2_O, C) N_2_O+N_2_, and D) the product ratio of N_2_ and N_2_O of the two land uses N_2_O+N_2_. Different letters in A-D indicate significant differences as derived from an LSD test (n=3, *p* value *<*0.05) comparing all groups of land use types and moisture contents. In addition, we report significant effects of the land use (grassland vs. cropland) and of soil moisture (pressure head of -160 cm vs. -40 cm vs -8 cm pressure head) and marked by ns, *, **, and *** for *p* values *>*0.05, *<*0.05, *<*0.01, and *<*0.001, respectively.

However, the proportion of N_2_O derived from denitrification (*Fp_N_*_2_*_O_*) consistently exceeded 0.6, approaching values close to one and unity at -8 cm pressure head (Fig. S3b). The calculated enrichment of ^15^N in the nitrate pool undergoing denitrification (ap-values) was significantly greater in the grassland across all moisture levels (Fig. S3a). This corroborates the trends observed in the measured ^15^N values of extracted nitrate (Fig. S3D) and ammonium (Fig. S3E), which aligned well with these enrichment data (Fig. S3G). Together with the high ^15^*N − NO*_3_^-^ values in all samples, this agreement in the data underlines the homogeneity of the ^15^N labeling as it shows that ^15^N of the denitrification hotspots (^15^N enrichment of NO_3_^-^ in anoxic microsites, ap-values), were identical with the ^15^N enrichment of extracted bulk soil nitrate. NO_3_^-^ concentrations after incubation did not significantly vary by land use or moisture conditions (Fig. S3C). Yet, the numerically lowest NO_3_^-^ concentrations were noted under the wettest conditions tested, aligning with the observed increase in N_2_O+N_2_ fluxes. The determined NH_4_^+^ concentration in none of the samples exceeded the quantification limit of 0.02 mmol/L, roughly corresponding to *<*5 µg/g soil.

### 3.2 Explanatory variables derived from X-ray CT

#### 3.2.1 Distance to air-filled pores

Sensitivity analyses revealed threshold distances almost identical for the cropland and the grassland (Table 1, 0.9 mm and 1.02 mm, respectively), which were used to calculate the potential anaerobic volume, i.e. the soil volume to far away from air-filled pores to maintain sufficient diffusive oxygen supply (Table 1). Note that the p-value for the cropland was the lowest for all distances at 0.9 mm, yet not significant (p*>*0.05). Under dry conditions, in which the majority of macropores are air-filled (Table S1), occurrences of distances to air-filled pores exceeding 1 mm were nearly nonexistent (Fig. S4). As the moisture content increases, more and more pores fill, thereby extending the distances to the air-filled pores (Fig. S4). Correspondingly, the potential anaerobic soil volume exhibited an increasing trend from the driest to the wettest soil, with numerically higher values observed in the cropland (Fig. 3b) caused by distinct differences in the pore structure between the two land uses, i.e. by the higher share of large distances (compacted soil areas) in the cropland soil (Fig. S4). However, notable outliers were observed in the grassland soil, with some cores displaying high potential anaerobic volumes, emphasizing the heterogeneity in the field. In the case of the grassland, there was a strong association between the potential anaerobic volume and the N_2_O+N_2_ fluxes (Fig. 3a, R^2^=0.82, *p<*0.001). For the cropland, however, even samples with relatively high potential anaerobic soil volumes resulted in fluxes of N_2_O+N_2_ close to 0. Note that the potential anaerobic volume is based on 3D pore distances only and is therefore a proxy for O_2_ supply without information about the heterogeneous distribution of O_2_ demand.

**Figure 3.**
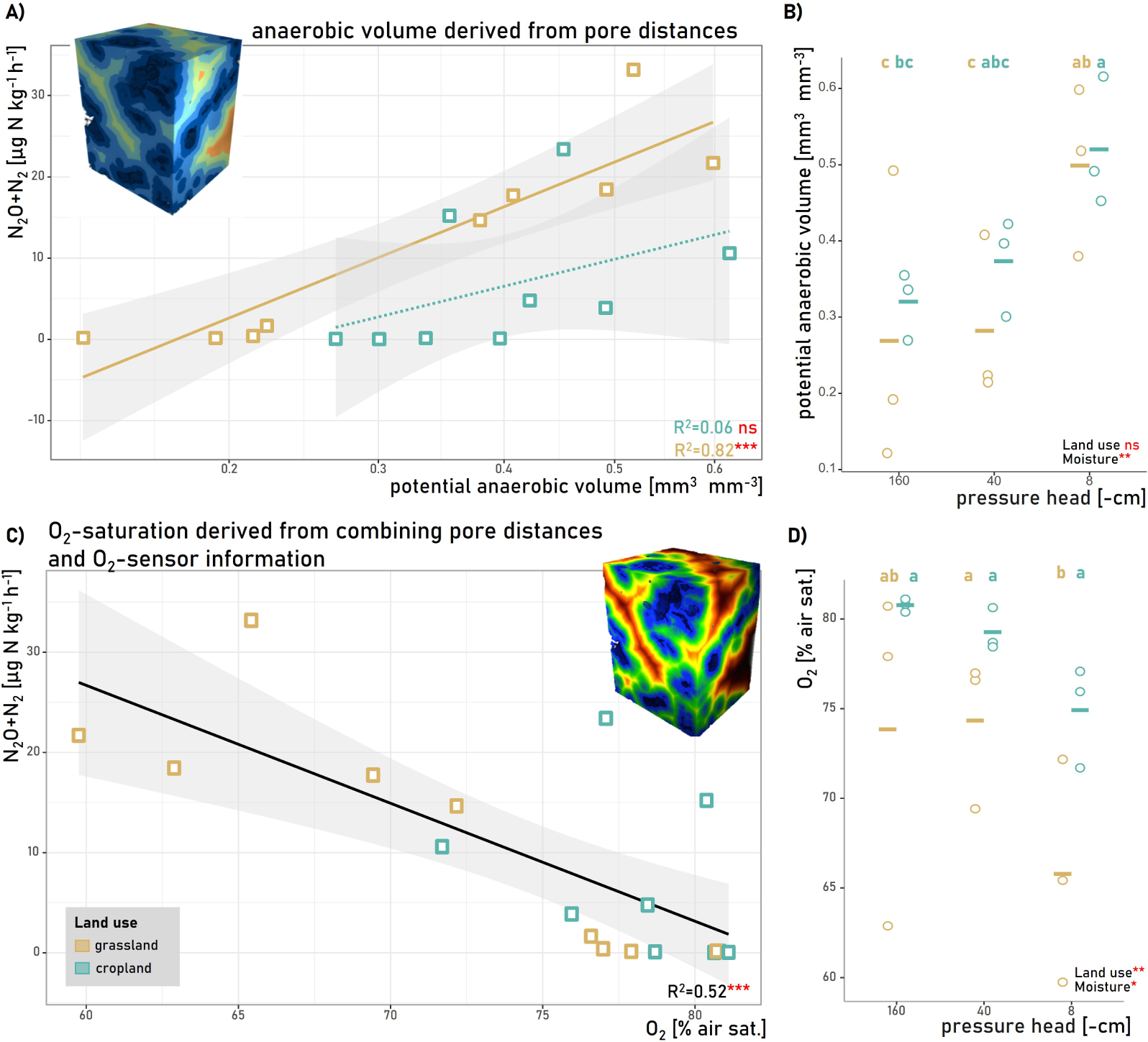
The anaerobic volume as a driver of denitrification. A) Association of the potential anaerobic soil volume (derived from X-ray CT) with N_2_O+N_2_ and B) respective mean volumes of a soil core (circles) and the mean of the treatment (crossbar) for the two different land uses and the three investigated water potentials of the potential anaerobic soil volume. C) Association of the mean O_2_-concentration in % of air saturation of the soil core (derived by a combination of X-ray CT and microsensors) with N_2_O+N_2_ fluxes and D) respective mean O_2_-concentration of a soil core (circles) and the mean of the treatment (crossbar) for the potential anaerobic soil volume. The solid lines in A) and C) represent fitted linear models. Respective R^2^ and significance level are given in the corner of the graphs. Different letters in B and D indicate significant differences as derived from an LSD test comparing all groups of land use types and moisture contents (n=3, *p* value *<*0.05). In addition, we report significant effects of the land use (grassland vs. cropland) and of soil moisture (pressure head of -160 cm vs. -40 cm vs -8 cm) marked by ns, *, **, and *** for *p*-values *>*0.05, *<*0.05, *<*0.01, and *<*0.001, respectively.

**Table 1.** Mean values and standard deviation for basic parameters of the two land uses under field structured conditions in comparison to sieved soil as presented in Rohe et al. (2021) (n=3 moisture contents). The distance threshold describes the minimum distance to air-filled pores above which the soil was denoted as anaerobic and is the result of linear regression analysis of the association of N_2_O+N_2_ with X-ray CT-derived information.

|  | grassland |  | cropland |  |
| --- | --- | --- | --- | --- |
|  | repacked | structured | repacked | structured |
| distance threshold [mm] | 5 | 1.02 | 5 | 0.9 |
| potential anaerobic volume [$\text{cm}^3 \text{ cm}^{-3}$] | $0.32 \pm 0.42a$ | $0.35 \pm 0.13a$ | $0.31 \pm 0.37a$ | $0.40 \pm 0.1a$ |
| $\text{O}_2$ [% air-sat.] | $48.97 \pm 13.55b$ | $72.28 \pm 20.78a$ | $48.1 \pm 15.98b$ | $75.73 \pm 11.71a$ |
| $\text{N}_2\text{O}$ [$\mu\text{g N kg}^{-1} \text{ h}^{-1}$] | $11.37 \pm 9.75a$ | $8.01 \pm 8.54ab$ | $3.37 \pm 2.88bc$ | $2.44 \pm 3.1c$ |
| $\text{CO}_2$ [$\mu\text{g C kg}^{-1} \text{ h}^{-1}$] | $435.29 \pm 203.62a$ | $433.29 \pm 142.84a$ | $126.67 \pm 53.94b$ | $182.49 \pm 79.24b$ |
| $\text{N}_2\text{O}+\text{N}_2$ [$\mu\text{g N kg}^{-1} \text{ h}^{-1}$] | $18.21 \pm 12.57a$ | $12.01 \pm 11.97ab$ | $5.75 \pm 5.12b$ | $6.46 \pm 8.32b$ |
| $\text{N}_2\text{O} / (\text{N}_2\text{O}+\text{N}_2)$ | $0.57 \pm 0.23a$ | $0.6 \pm 0.13a$ | $0.45 \pm 0.3b$ | $0.4 \pm 0.21b$ |

#### 3.2.2 3D oxygen distribution

In addition to monitoring greenhouse gas fluxes, we tracked O_2_ in % air saturation using microsensors throughout the experiment (Fig. S5). The majority of these microsensors consistently indicated air saturation levels close to 100% throughout the entire incubation period, with only a few registering values below 50% at the onset of the experiment and thereafter. Under wetter conditions (-8 cm), there was a collective decrease in O_2_ readings over time, although saturation levels never approached zero. The O_2_ saturation at the end of the incubation displayed a significant correlation with the distance to air-filled pores as computed for the sensor tips (Fig. S7), albeit with some outliers showing relatively low O_2_ saturation at shorter pore distances.

Based on these associations, we derived the 3D distribution of O_2_ (Fig. 1) along with corresponding average values (Fig. 3d). In comparison to the potential anaerobic soil volume, the average O_2_ saturation was less affected by soil moisture, and contrary to expectations based on the potential anaerobic soil volumes, the cropland exhibited higher O_2_ saturation levels compared to the grassland (78.3% in the cropland and 71.3% in the grassland). Additionally, the average 3D O_2_ saturation displayed a strong negative correlation with N_2_O+N_2_ fluxes (R^2^=0.52, *p<*0.001) and was unaffected by land use type (Fig. 3c).

#### 3.2.3 Position of POM

We enhanced the resolution of X-ray CT scanning by subsampling the incubated cores, enabling the segmentation of POM (Fig. 1). Consistent with the higher total carbon concentration in grassland, we observed a significantly higher share of POM in the grassland (4.0-4.6%) compared to the cropland with values ranging between 1. and 2.7% (Table S1). The distribution of POM was strongly skewed towards low distances to air-filled pores, with nearly all POM situated within or close to air-filled pores (Fig. S4a). Thus, for both land uses, the mean air distance of the POM in relation to air distances of the soil matrix was *<*0.1. However, this distribution indicator revealed consistently higher values in the grassland than in the cropland indicating a better incorporation of POM into the soil matrix within the grassland (grassland: 0.031 - 0.045, cropland: 0.01 - 0.02, Fig. S4b). However, as moisture content increased and some macropores containing POM filled with water (Table S1), distances of POM to pores increased relative to the matrix distances. Interestingly, the portion of POM identified as intact root segments exhibited a similar pattern, with most roots located within or close to air-filled pores (Fig. S6). The volume of proximal POM, defined as POM closer than 0.59 mm to air-filled pores as indicated by sensitivity analysis related to N_2_O+N_2_ fluxes, showed no significant variation with soil moisture but was higher in grassland than in cropland (Fig. 4c, 0.04 mm^3^ mm^-3^ compared to 0.01 mm^3^ mm^-3^). On the other hand, distal POM, increased with moisture content and was higher in the grassland across all moisture conditions (Fig. 4f). While proximal POM did not show a significant correlation with N_2_O+N_2_ fluxes from denitrification, it was strongly associated with CO_2_ fluxes (R^2^=0.7, *p* value*<*0.001, Fig. 4a,b). In contrast, distal POM displayed the opposite pattern, with a strong association with N_2_O+N_2_ fluxes produced by denitrification (R^2^=0.79, *p* value*<*0.001) and no association with CO_2_ fluxes (Fig. 4d,e). The association of the distal POM with N_2_O alone was even stronger (R^2^=0.90, *p* value *<*0.001, Fig. S8) compared to N_2_O+N_2_. Additionally, the amount of distal POM displayed a clear negative correlation with NO_3_^-^ concentrations, in accordance with its link to denitrification processes (Fig. S3f).

**Figure 4.**
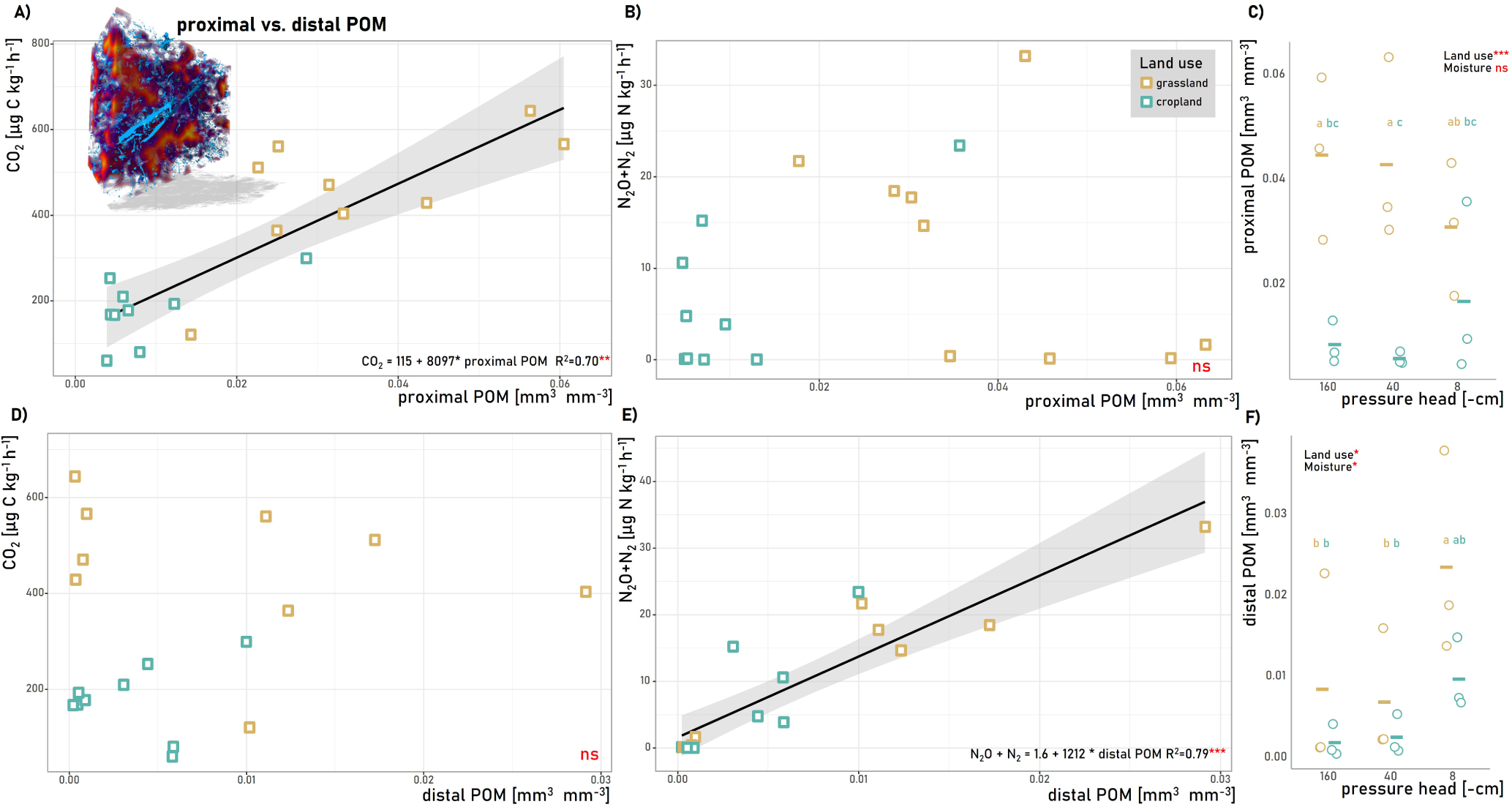
The position of POM as the driver of denitrification. Association of the amount of proximal POM (to air-filled pores) and CO_2_ fluxes (A), N_2_O+N_2_ fluxes produced through denitrification (B), and respective mean values of the amount of proximal POM volume (C) for the two different land uses and the three investigated water potentials. Association of the amount of proximal POM as well as distal POM with CO_2_ fluxes (A, C), N_2_O+N_2_ fluxes (B, E), and respective mean values of the amount of the proximal POM volume (C), and the distal POM volume (F) for the two different land uses and the three investigated water potentials. The solid lines in A), and E) represent fitted linear models. Note that the linear relationships in B) and D) were non-significant and thus lines are not shown. Respective R^2^ and significance level are given in the corner of the graphs. Different letters in B and D indicate significant differences as derived from an LSD test comparing all groups of land use types and moisture contents (n=3, *p* value *<*0.05). In addition, we report significant effects of the land use (grassland vs. cropland) and of soil moisture (pressure head of -160 cm vs. -40 cm vs -8 cm) marked by ns, *, **, and *** for *p*-values *>*0.05, *<*0.05, *<*0.01, and *<*0.001, respectively.

#### 3.2.4 CO_2_ equivalents

We found compelling associations between the POM volume and GHG production quantified as the CO_2_-equivalents of CO_2_ and N_2_O combined (Fig. 5a). Note that CH_4_-production was below the detection limit. Distal POM demonstrated a steeper increase in association with GHG production, reflecting its role in the heightened production of the more potent greenhouse gas N_2_O. Furthermore, GHG production displayed significant associations with both land use type and moisture content. The grassland consistently exhibited numerically higher GHG production compared to the cropland across all moisture conditions (5.26 mg CO_2_-eq kg^-1^ h^-1^ in the grassland and 1.79 mg CO_2_-eq kg^-1^ h^-1^ in the cropland). This result aligns with the above findings regarding the association of proximal POM with CO_2_ and distal POM with N_2_O (Fig. 3), leading to the highest CO_2_-equivalents in the wettest grassland soil cores when most of the POM becomes distal.

**Figure 5.**
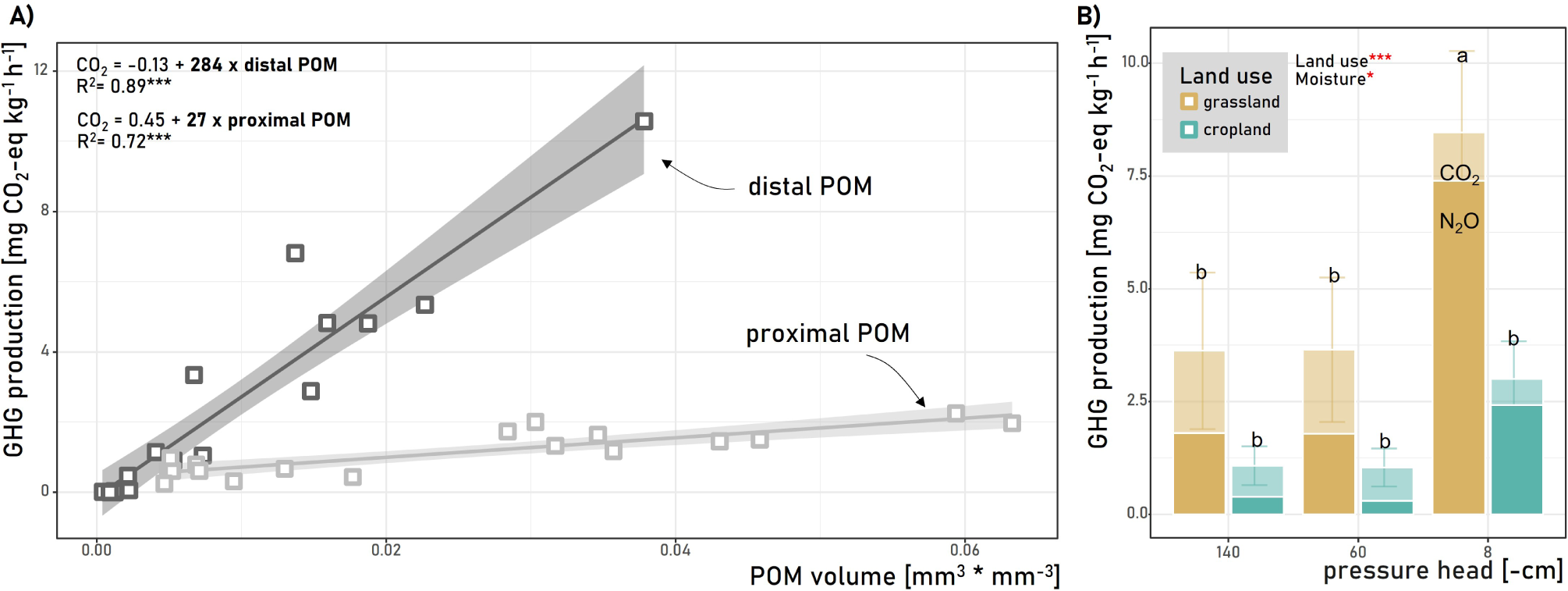
Impact of the share of proximal POM and distal POM (to air-filled pores) on the production of GHG (CO_2_ equivalents of CO_2_ and N_2_O) (A) and respective mean values and standard deviations for the two different land uses and the three investigated water potentials. The solid lines represent fitted linear models. Respective R_2_ and significance level are given in the corner of the graphs. Different letters in B indicate significant differences as derived from an LSD test comparing all groups of land use types and moisture contents (n=3, *p*-value *<*0.05). In addition, we report significant effects of the land use (grassland vs. cropland) and of soil moisture (pressure head of -160 cm vs. -40 cm vs -8 cm) marked by ns, *, **, and *** for *p*-values *>*0.05, *<*0.05, *<*0.01, and *<*0.001, respectively.

#### 3.2.5 Comparison to sieved soil

The comparison between repacked soil cores used in Rohe, Apelt, et al. (2021) and the structured soil cores in our study revealed notable differences in the estimated distance thresholds used to calculate the potential anaerobic volume, with much lower distance thresholds in the structured cores (Table 1). Surprisingly, despite these differences, the potential anaerobic volume averaged across all three moisture contents did not show significant differences between the sieved and structured versions of the soil. However, significant disparities emerged in the O_2_-saturation calculated from the microsensors installed. In the repacked soil, grassland cores exhibited a mean O_2_ saturation of 49.0±13.6%, and cropland cores showed 48.1±15.98%. In contrast, the structured cores displayed significantly higher O_2_ saturation, with grassland at 72.3±20.8% and cropland at 75.7±11.7%. Despite the higher O_2_ saturation in structured cores, N_2_O, N_2_O+N_2_, CO_2_ fluxes, and the product ratio of N_2_O and N_2_ were consistently higher in the grassland compared to cores from the cropland, regardless of whether they were sieved or structured.

## 4 DISCUSSION

In this study, we investigated the dynamics of CO_2_, N_2_, and N_2_O emissions, in soil cores under varying moisture conditions and land use types. Utilizing a combination of gas flux measurements, microsensor data, and X-ray CT scanning, we sought to understand the underlying mechanisms driving the production of these gases and their relationship with soil properties. Our findings revealed significant variability in CO_2_, N_2_ and N_2_O emissions and in the fraction of N_2_O from denitrification across different land use types and moisture levels (Fig. 2). Moreover, our investigation into POM distribution highlighted its influence on GHG production, with both proximal and distal POM showing associations with CO_2_, N_2_ and N_2_O fluxes, respectively (Fig. 5).

### 4.1 Denitrification induced by POM

Similar to previous studies, our analysis of the 3D O_2_ distribution demonstrated the direct and decisive influence of O_2_ on N_2_O in structured soils (Du et al., 2023; Song et al., 2019; Wei et al., 2023, Fig. 3). In addition to these studies, however, we also analyzed N_2_ to get an estimate of the complete potential of N losses through denitrification. The construction of the 3D O_2_ distribution was mainly based on structural properties, but not on hotspots of POM. The predictive power of the calculated O_2_ concentrations in the columns is therefore limited (Fig. 3) as it does not capture outliers with low concentrations at comparably small distances (Fig. S7), which are likely a result of hotspots. The extent of denitrification in soils is a result of balancing between O_2_ demand and supply (Robertson, 2023; Rohe, Apelt, et al., 2021). The O_2_ supply, on the one side, is controlled by the soil structure through the regulation of diffusion distances and the establishment of potential anaerobic volumes (Du et al., 2023; Lucas, Gil, et al., 2023; Rohe, Apelt, et al., 2021, Fig. 3). Optimal soil moisture conditions for intensive N_2_O emissions typically fall within a broad range of around 70% water-filled pore space (WFPS), with variations observed among different land uses (Bateman & Baggs, 2005; Rohe, Apelt, et al., 2021; Ruser et al., 2006; Weier et al., 1993). This variability can be attributed to the emergence of anaerobic soil volumes in hotspots of high oxygen consumption within an otherwise well-aerated soil. Although it’s known for decades, that such hotspots are important drivers of denitrification as they control the other side of the O_2_ balance, i.e. the O_2_ demand (Parkin, 1987), this work emphasizes the need to precisely locate these hotspots to gain a better understanding of the denitrification potential. For the first time, we were able to prove that the position of POM within the heterogeneous pore system, or more precisely its distance to air-filled pores, is a decisive factor for balancing between elevated CO_2_ and N_2_O+N_2_ emissions from hotspots in soils. While for POM close or within air-filled pores the enhanced O_2_ demand can mostly be satisfied and predominately CO_2_ is produced, it is the anaerobic microsite formed on distal POM, where all conditions necessary for denitrification, i.e. low O_2_, high C, and N are met (Fig. 4). This is in agreement with findings revealing that large mean distances of POM to air-filled pores are associated with N_2_O emissions (Lucas, Santiago, et al., 2023; Ortega-Ramírez et al., 2023). The significance of POM as a hotspot for denitrification is further highlighted through the comparison with the sieved soil studied in Rohe, Apelt, et al. (2021). Our findings reveal a stark contrast in the threshold for oxygen limitations, which was observed to exceed 5 mm in sieved soils lacking POM, as opposed to the approximately 1 mm threshold in naturally structured field soils (Table 1). This discrepancy underscores the limited capacity for denitrification initiation within soil matrices devoid of POM hotspots (Schlüter et al., 2024), given the need for diffusion length *>*5 mm for the relatively low O_2_ demand. These long distances are scarce in structured field soils even under conditions of low water content (Fig. S4a). Last, the role of distal POM (to air-filled pores) in the fate of NO_3_^-^ is supported by the observed correlation between areas with low NO_3_ concentrations and those with high levels of distal POM (Fig. S3F). We determined a fixed threshold to differentiate between proximal and distal POM from X-ray CT images taken at the end of the incubation (Table 1), assuming a constant O_2_ demand for all POM hotspots. However, O_2_ (Fig. S1) and N_2_O+N_2_ (Fig. S2) levels were not constant during incubation. This is due to 1. a delay until anoxic centers are fully developed and all denitrification enzymes are expressed, 2. a decrease in N_2_O, CO_2_, and O_2_ over time because carbon substrates and nitrate are consumed and 3. internal water reconfiguration that changes air continuity and thus local O_2_ concentrations. In addition, under field conditions, continuous inputs of fresh labile carbon from living plants or organic fertilizers may significantly modify the distance threshold, as revealed in a recent meta-analysis (Schlüter et al., 2024). Our incubation study did not replicate this dynamic nature, highlighting a need for future experiments to address this variability. However, the analysis by Schlüter et al. (2024), found a median distance threshold of 0.9 mm for labile carbon mixed into the soil, which aligns closely with the thresholds computed in this study (Table 1). In conclusion, to refine predictions of denitrification processes, future models should include POM as a dynamic state variable and parameterize the physical separation of POM from air-filled pores as a function of water content. Note that the linear relationship between distal POM and N_2_O+N_2_ produced by denitrification found in this study was unaffected by land use. This holds promise for the development of a universal empirical equation and could significantly enhance the predictive accuracy of denitrification models. Note that the traditional definitions of free and occluded POM, while often defined operationally by the fractionation method (Leuthold et al., 2023; Wagai et al., 2009), are similar to the here-defined proximal and distal POM. Specifically, free POM is believed to be found in macropores between soil aggregates, while occluded POM is embedded within these aggregates, providing it with a degree of protection (Brodowski et al., 2006; Golchin et al., 1994; Kölbl et al., 2005). However, the CT-based identification of particulate organic matter (POM) fractions marks a significant advancement over the traditional density fractionation method for categorizing occluded and free POM as hotspots of denitrification and aerobic respiration, respectively. This is due to the dynamic nature of CT-derived POM fractions of distal and proximal POM, whose proportions fluctuate with changes in water content, as illustrated in Figure 4c. This fluctuation enables a more accurate prediction of denitrification by capturing the interplay between oxygen demand and supply, under varying moisture levels. Consequently, there is a compelling case for further investigation to develop comparably effective denitrification predictors that integrate the methodological simpler and often applied occluded POM determination through fractionation with pore structural measures, such as pore size distribution. At the same time, the mechanisms that lead to a different POM distribution in the pore space must be further investigated to improve future denitrification models taking into account the heterogeneity on the pore scale for field scale prediction.

### 4.2 Denitrification in cropland vs. grassland

The investigated grassland and cropland featured different pore structures and POM abundances (Fig. 6). Notably, despite larger potential anaerobic volumes brought about by pore structure (3b), the cropland had higher O_2_ saturation and lower denitrification activity because of lower POM content in general, and lower distal POM (to air-filled pores) content in particular (Table 1). Thus, the position of POM in the cropland, often near or within pores, limited denitrification rates despite the dense structure. This configuration resulted in relatively low denitrification rates compared to the grassland, where the abundance of distal POM was notably higher (Fig. 4 and Fig. 5b). One reason for this non-random distribution is that a large share of POM originates from roots that predominately grow into looser areas or directly into macropores (Lucas et al., 2019a, 2022; White & Kirkegaard, 2010, Fig. S6). and thus POM fragments are typically located close to air-filled macropores (Kravchenko et al., 2018; Schlüter et al., 2022). Conversely, the grassland, characterized by its lower bulk density (Table 1) and higher SOC content, presented a contrasting scenario with reduced potential anaerobic volumes and consequently enhanced O_2_ supply, particularly under dry conditions (Fig. 3). However, the advantage in O_2_ supply was offset by the higher O_2_ demand attributed to the substantial amounts of POM present in the grassland (Table S1). Under dry conditions, microbial respiration primarily drove the utilization of this POM, especially within or proximate to macropores. Thus, the grassland exhibited higher CO_2_ fluxes, while under drier conditions N_2_O fluxes were similar to those in the cropland. As moisture levels increased and macropores became increasingly saturated, the O_2_ supply to microbial hotspots was reduced. Consequently, the accumulation of large amounts of distal POM under moist conditions triggered N_2_O emissions and consequently the overall GHG production (Fig. 5). While tillage practices on croplands lead to vertical mixing of crop residues with soil, it also reduces the amount of soil fauna like anecic earthworms, which were shown to incorporate POM into the soil matrix (Briones & Schmidt, 2017; Leuther et al., 2023; Wilkinson et al., 2009; Yang & Wander, 1999). In contrast, on the grassland, permanent vegetation cover exhibits higher bioturbation rates, resulting in greater incorporation of POM into the soil matrix (Kraus et al., 2022; Stolze et al., 2022). and a more even POM distribution (Fig. S4b). The more evenly distributed POM in the grassland compared to the cropland led to the high association of N_2_O+N_2_ emissions with CT-derived potential anaerobic soil volumes (Fig. 3). This indicates that in soils with an even distribution of hotspots (oxygen demand), the heterogeneity in denitrification is mainly driven by variations in oxygen supply at the microscale (Lucas, Gil, et al., 2023; Rohe, Apelt, et al., 2021). However, our observations underline the major importance of denitrification for the total N_2_O emissions from soil, particularly under near saturated conditions when N_2_O production rates were high and almost all N_2_O is produced by denitrification (Fig. S3b), corroborating previous findings (Gao et al., 2023; Liang & Robertson, 2021; Lucas, Gil, et al., 2023; Ostrom et al., 2021). In addition, the sustained N_2_O emissions observed even after 10 days of incubation support the notion, that hotpots of N_2_O can persist for several weeks under static conditions (Christensen et al., 1990). However, denitrification completeness, often expressed by the product stoichiometry, N_2_O/(N_2_O+N_2_), can vary a lot in soils depending on environmental factors. Dhaliwal et al. (2024) and Rohe, Apelt, et al. (2021) suggested that low macroporosity and thus large diffusion distances of N_2_O through the soil matrix favor N_2_O reduction. However, the structured cores used in this study had the same product ratio of N_2_O and N_2_ compared to the sieved soils used in Rohe, Apelt, et al. (2021) despite much higher diffusion distances (Table 1). This suggests that diffusion distance does not exclusively control this ratio. The disparity in soil pH levels offers a plausible explanation for the observed differences in product ratios (Fig. 2d) among the land uses examined (Čuhel & Šimek, 2011). Yet, under the wettest conditions, both land use types exhibited similarly high product ratios (*>*0.6) and thus higher N_2_O fluxes compared to N_2_ at high denitrification rates - this phenomenon, although noted before (Kemmann et al., 2021; Lucas, Santiago, et al., 2023), is controversial to the basic understanding of denitrification, where decreased O_2_ concentration leads to an increase of N_2_O reductase (NOS) and thus to more N_2_ (Morley & Baggs, 2010; Robertson & Groffman, 2024). A potential explanation, awaiting experimental validation, emerges from observations of the model denitrifier Paracoccus denitrificans, which under hypoxic conditions synthesizes NOS in all cells, whereas other enzymes in the denitrification pathway are scarcely expressed (Lycus et al., 2018). This bet-hedging strategy is thought to be common among denitrifying bacteria and may lead to NOS activity under moderate moisture conditions characterized by multiple oxic and hypoxic microsites (Lycus et al., 2018). Conversely, under the wettest conditions tested, anoxic zones become more prevalent, prompting a stronger expression of nitrate and nitrite reductases and therefore N_2_O emissions.

**Figure 6.**
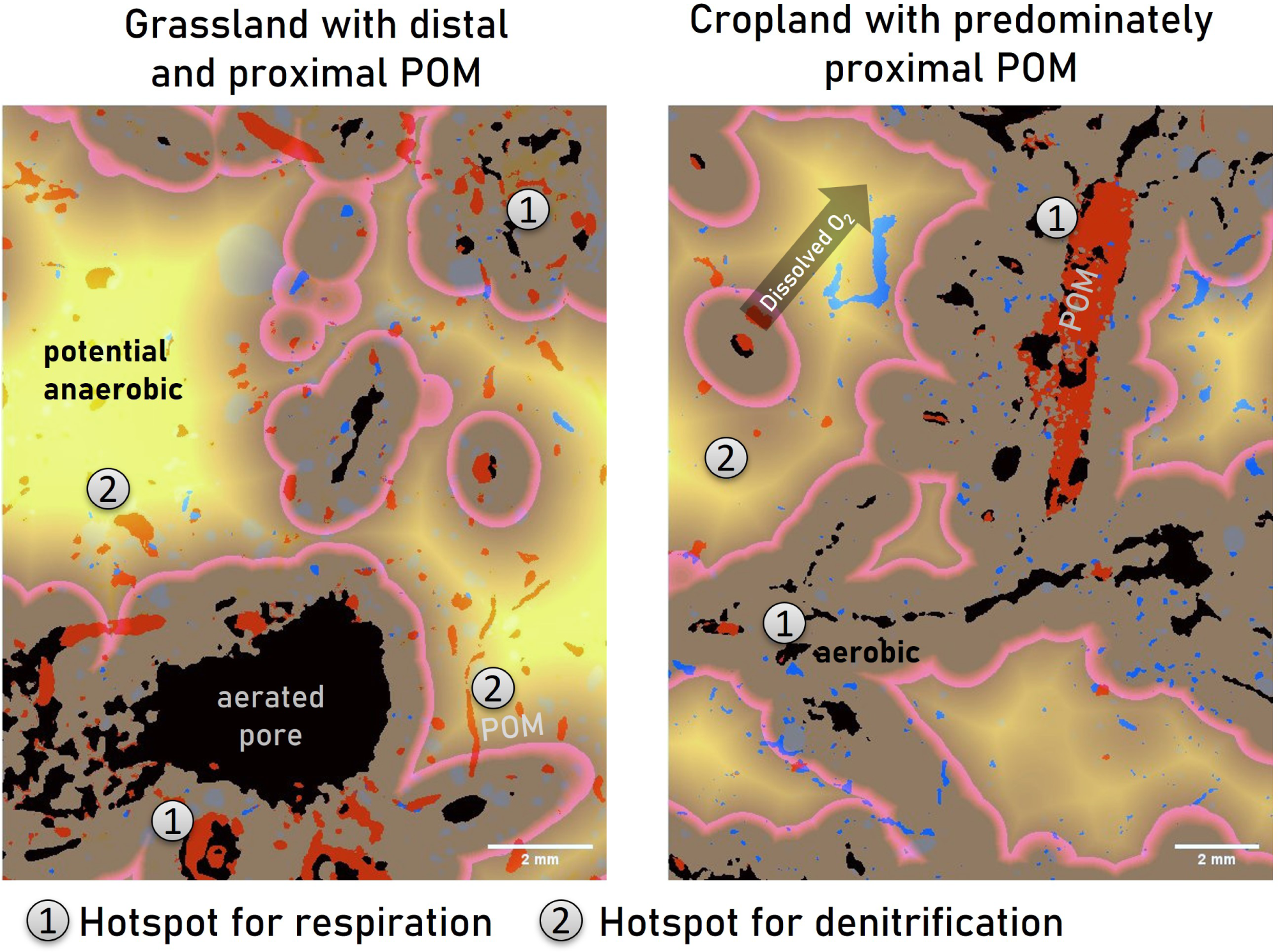
2D slices of segmented X-ray CT images of the grassland and the cropland. The distribution of POM (red) to air-filled pores (black) is decisive for the formation of hotspots of denitrification vs. respiration. Hotspots of respiration are created by POM at low distances to air-filled pores, while POM in the potential anaerobic volume (yellow area) leads to a hotspot of denitrification. Note that, in the grassland the distribution of POM was more homogeneous in comparison to the cropland. The cropland was characterized by the presence of large volumes of potentially anaerobic volumes, but relatively few hotspots (distal POM) for denitrification in it.

### 4.3 Concluding remarks

Our observations underscore the intricate interplay between soil pore structure, moisture dynamics, and POM distribution in modulating denitrification processes and greenhouse gas emissions. Moreover, they emphasize new possibilities in soil N management and mitigation strategies aimed at curbing greenhouse gas emissions from terrestrial ecosystems, which include strategies for changing the distribution of POM, by utilising plants with roots that reuse a higher proportion of pores (Lucas et al., 2022). Recent literature emphasizes the critical role of anoxic microsites in regulating CO_2_ emissions (Keiluweit et al., 2016; Lacroix et al., 2022, 2023). Keiluweit et al. (2017) underscored the importance of oxygen limitation in soil domains with restricted oxygen supply, such as aggregates or peds, in modulating carbon storage dynamics. These studies collectively suggest that anoxic microsites are crucial in governing carbon storage in terrestrial ecosystems. Moreover, a recent perspective by Angst et al. (2023) emphasizes the need to consider the site-specific relevance of POM in carbon sequestration. These perspectives suggest that while matrix-associated organic matter accumulation has traditionally been a focus of carbon sequestration efforts, the persistence and significance of POM, particularly in its occluded form, should not be overlooked. Our findings align with this perspective, indicating that distal POM in the soil matrix indeed is protected from microbial respiration and therefore decreases CO_2_ losses. However, the positive effects on the climate balance associated with measures to increase carbon storage in soils can be reduced or even negated by N_2_O emissions (Guenet et al., 2021). Indeed, our data suggest the drawback in the overall climatic balance when storing POM in the soil matrix for C protection. Note that our incubation setups involved both, high moisture status and NO_3_^-^-levels. These conditions may be met in heavily managed soils during high rain events. Therefore, as previously suggested by modelling results, such a risk of increased N_2_O emissions from management practices targeting C accumulation can be mitigated by adjusting N fertilisation rates (Qiu et al., 2009; You et al., 2024).

## AUTHOR CONTRIBUTIONS

ML and BA conducted the incubation experiment. RW analyzed the IRMS data. CFS conducted the SpinMas analysis. LR and BA conducted pre-experiments and optimized the incubation setup. SS, HJV, ML, RW, and LR conceived the experiment. ML did the combined data analysis. Funding acquisition by HJV and SS. All authors have seen and approved the final version submitted and were part of the writing process.

## ACKNOWLEDGMENTS

We gratefully acknowledge the valuable insights and constructive feedback provided by the anonymous reviewers, which have significantly improved the quality of this manuscript. This study was funded by the Deutsche Forschungsgemeinschaft through the research unit DFG-FOR 2337: Denitrification in Agricultural Soils: Integrated Control and Modeling at Various Scale (DASIM), grant number 270261188.

## DATA AVAILABILITY STATEMENT

The datasets generated for this study can be found under the DOI:10.48758/ufz.14621.

