## Supplementary File for "The distribution of particulate organic matter in the heterogeneous soil matrix - balancing between aerobic respiration and denitrification"

#### Supplementary Material

### 1 SUPPLEMENTARY DATA

#### 1.1 Figures

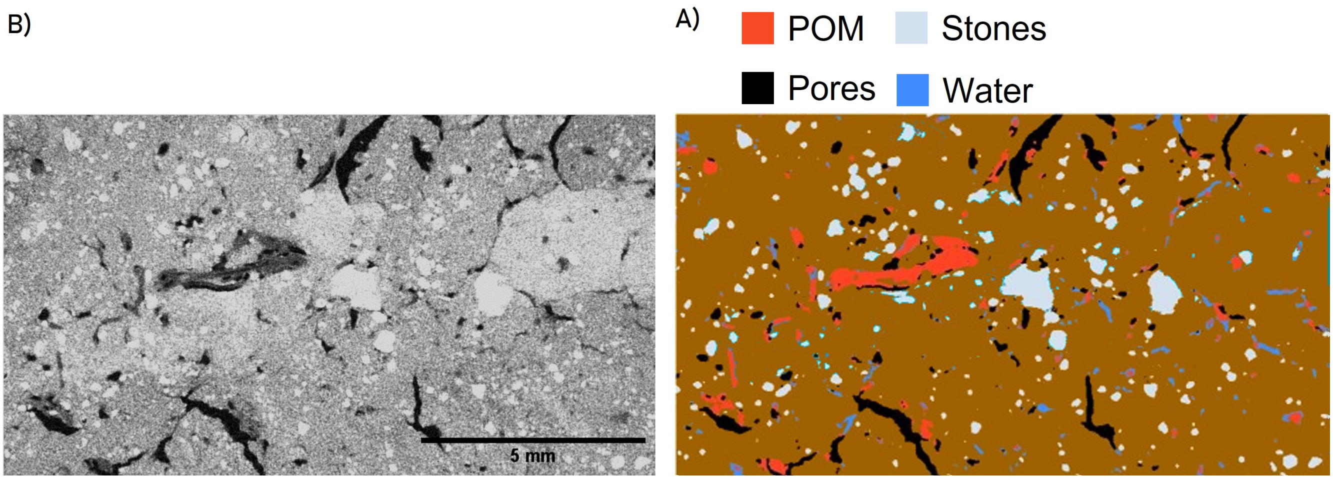


Figure S1: Slice of reconstructed, unfiltered X-ray CT image (A) and corresponding segmented image (B).

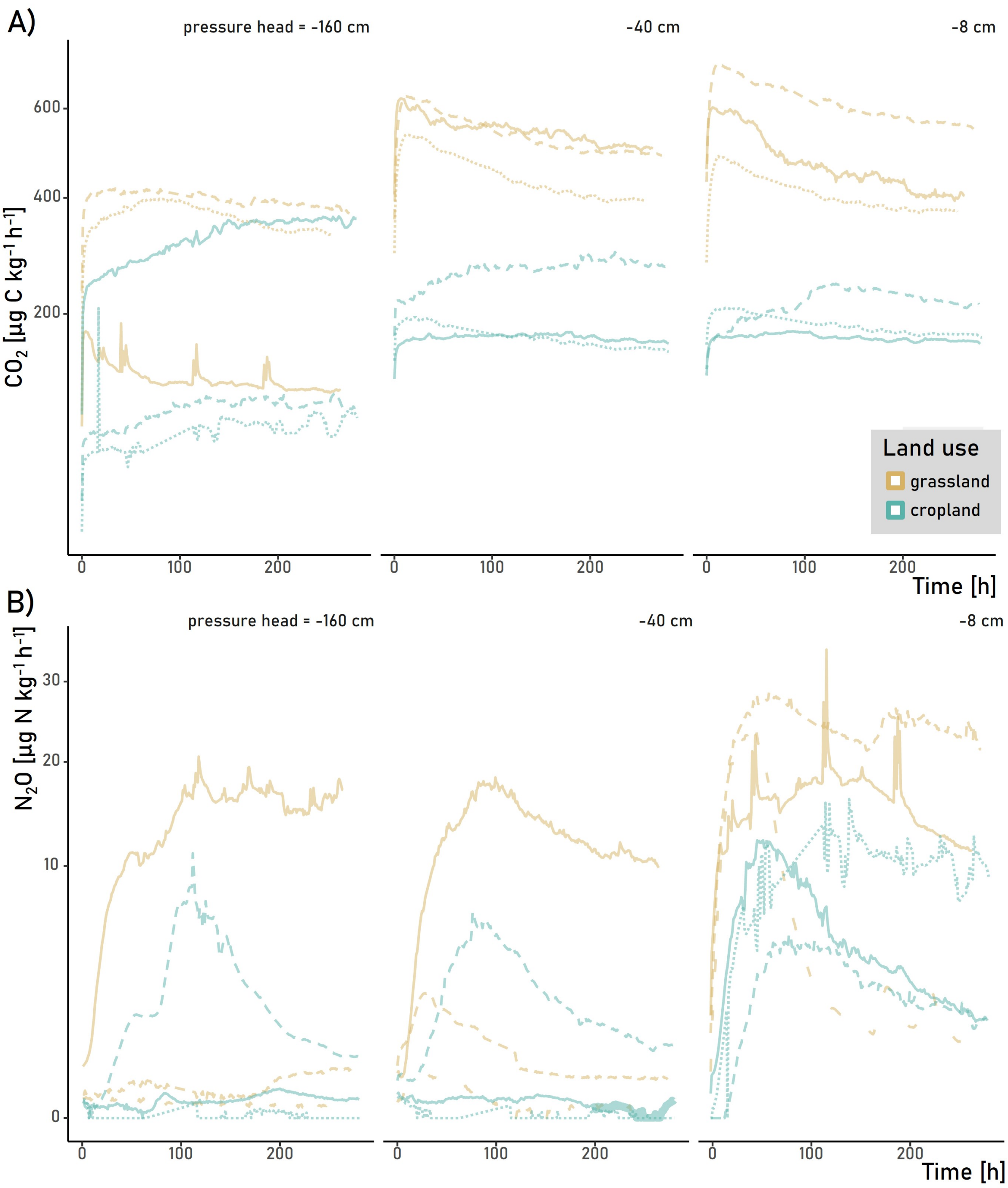


Figure S2: $CO_2$ (A) and $N_2O$ total (B) emissions during the incubation experiment at three different moisture contents. Each curve results from fluxes monitored hourly from one undisturbed soil core. Note that some of the large peaks are a result of pressure changes during sampling for IRMS measurements.

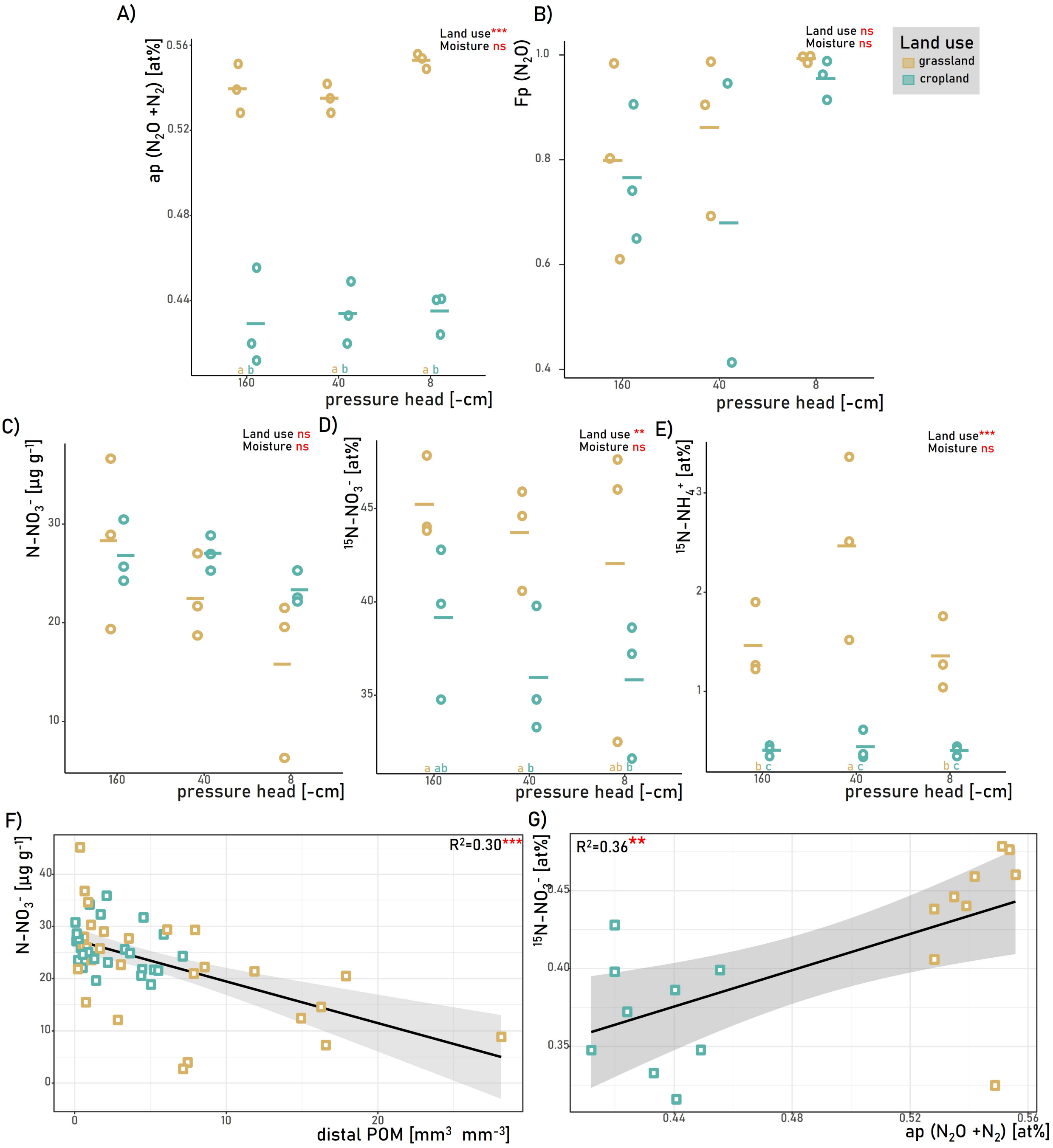


Figure S3: The distribution of the $^{15}$N-$NO_3^-$-label. A) Ap-values and B) Fp-values of the two land uses $N_2O$+$N_2$ at three different moisture contents. Note, that the ap-value is the calculated enrichment of the $NO_3^-$-pool denitrified to $N_2O$ and $N_2$. The Fp-value is the share of $N_2O$ derived from the labeled $NO_3^-$-pool, i.e. the share denitrification on the total $N_2O$ production. C) The amount of N-$NO_3^-$ at the end of the experiment as well as D) 15N-$NO_3^-$ and E) $^{15}$N-$NH_4^+$ at the end of the experiment within the two land uses and at the three investigated moisture contents. F) Association of distal POM and N-$NO_3^-$. Note that the distal POM was calculated for depths corresponding to three different locations of disturbed soil sampling. G) Association of the ap-values with $^{15}$N-$NO_3^-$. The solid lines in E) and F) represent fitted linear models. Respective $R^2$ and significance level are given in the corner of the graphs. Different letters in A-D indicate significant differences as derived from an LSD test comparing all groups of land use types and moisture contents (n=3, p-value <0.05). In addition, we report significant effects of the land use (grassland vs. cropland) and of soil moisture (matric potential of -160 cm vs. -40 cm vs -8 cm) marked by ns, *, **, and *** for p-values >0.05, <0.05, <0.01, and <0.001, respectively.

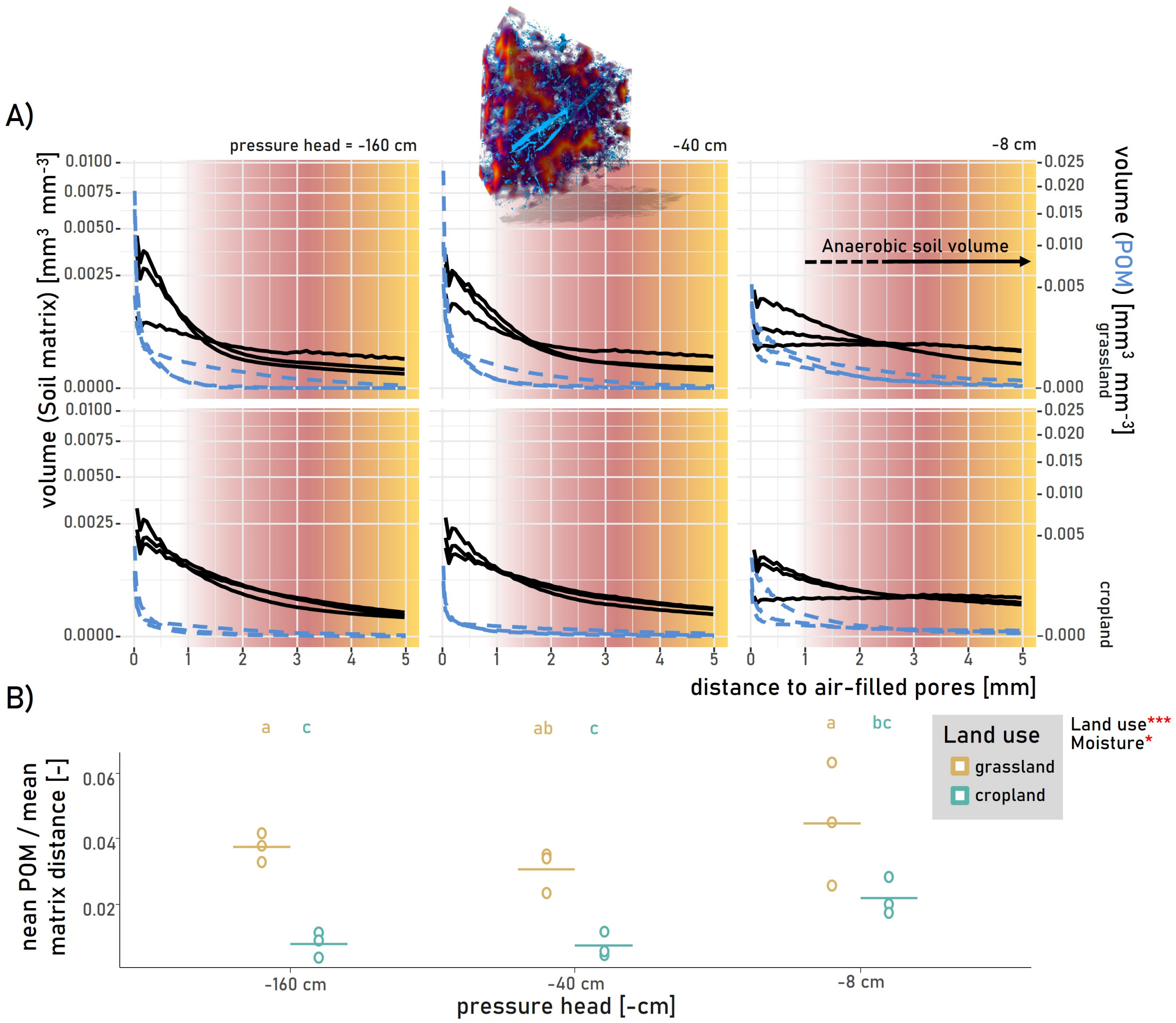


Figure S4: Histogram of distances A) and mean POM distance divided by mean matrix distance to air-filled pores B). The distances for every location within a soil column are shown to the next pore (black, A). Blue lines represent the distances of POM to the next pores. Background colors match the distances of the 3D visualization (upper left) and reveal the anaerobic soil volume fraction starting at distances of around 0.6 mm. Circles in B) represent the mean value of a core, while the straight lines are means per treatment for the two land uses at three different moisture contents In addition, we report significant effects of the land use (grassland vs. cropland) and of soil moisture (matric potential of -160 cm vs. -40 cm vs -8 cm) marked by ns, *, **, and *** for p-values >0.05, <0.05, <0.01, and <0.001, respectively.

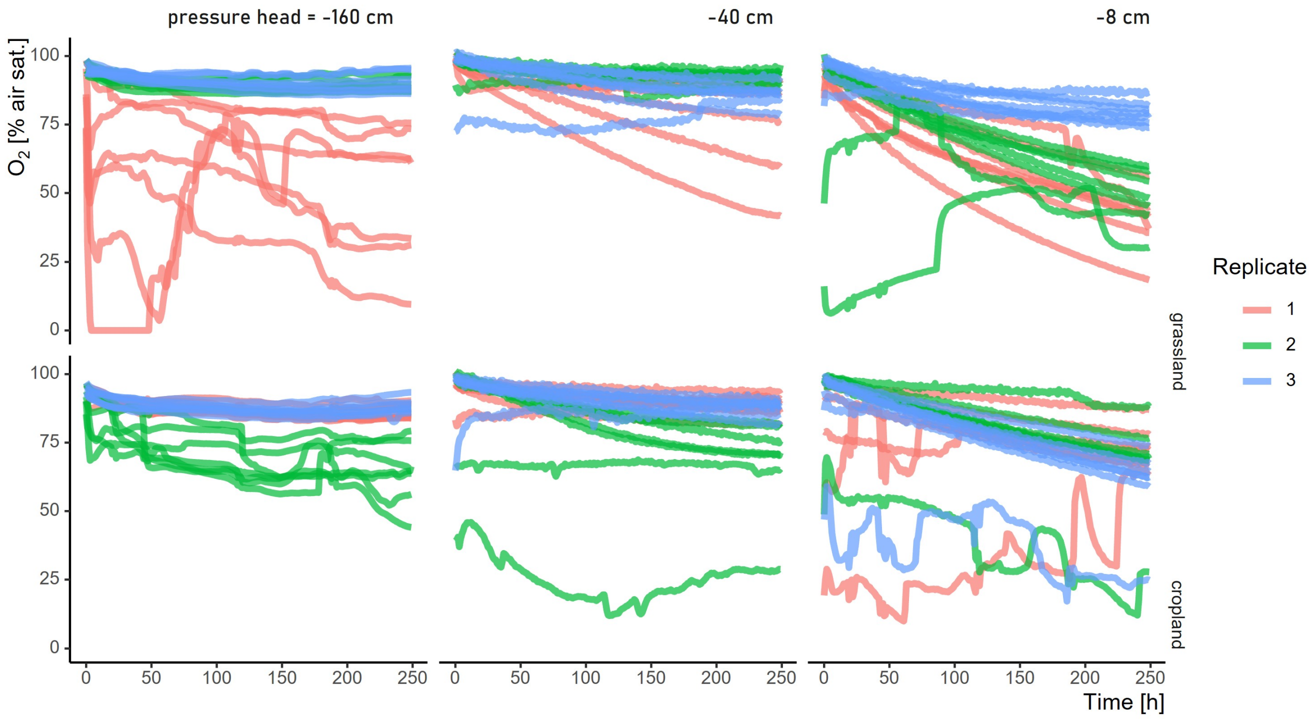


Figure S5: $O_2$-saturation during the incubation. Each line represents one out of seven $O_2$ microsensors installed in the replicated soil cores under three different soil moisture contents. $O_2$ concentration is given as % relative to saturated air and was logged twice an hour.

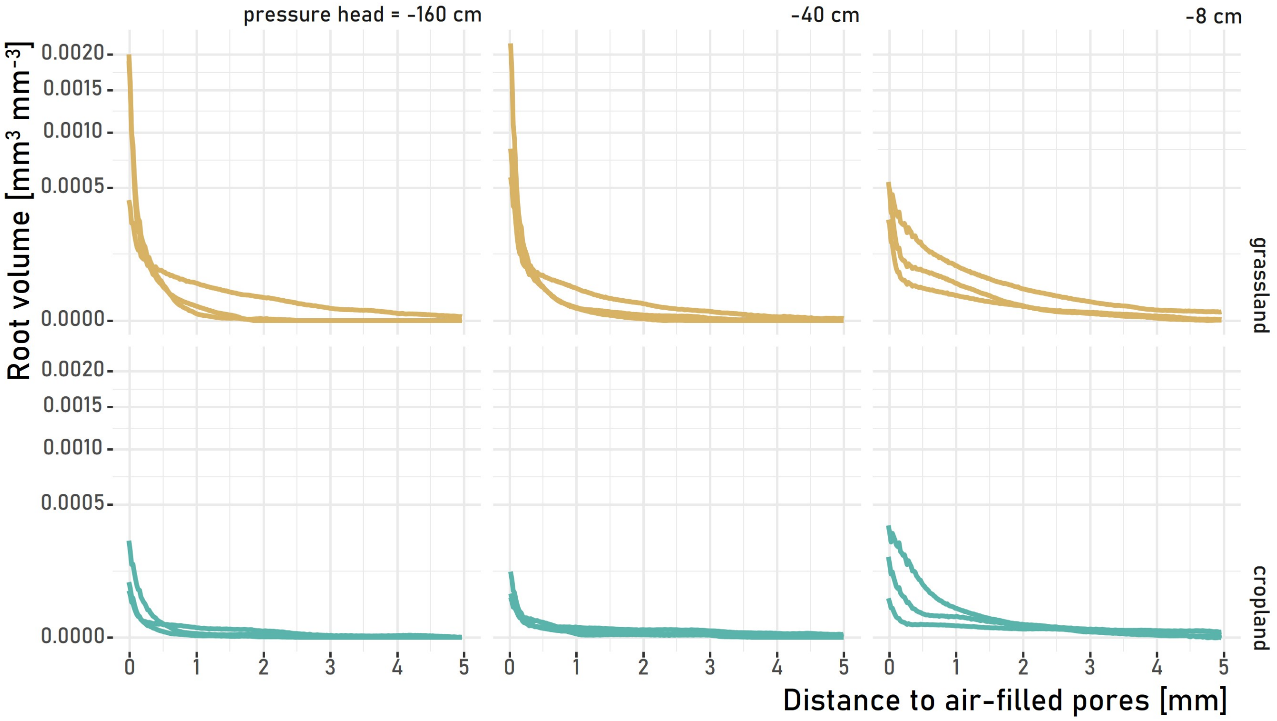


Figure S6: Histogram of distances from intact root segments. The Euclidean distances for each root voxel within a soil column to the next air-filled pore at the three moisture contents are shown.

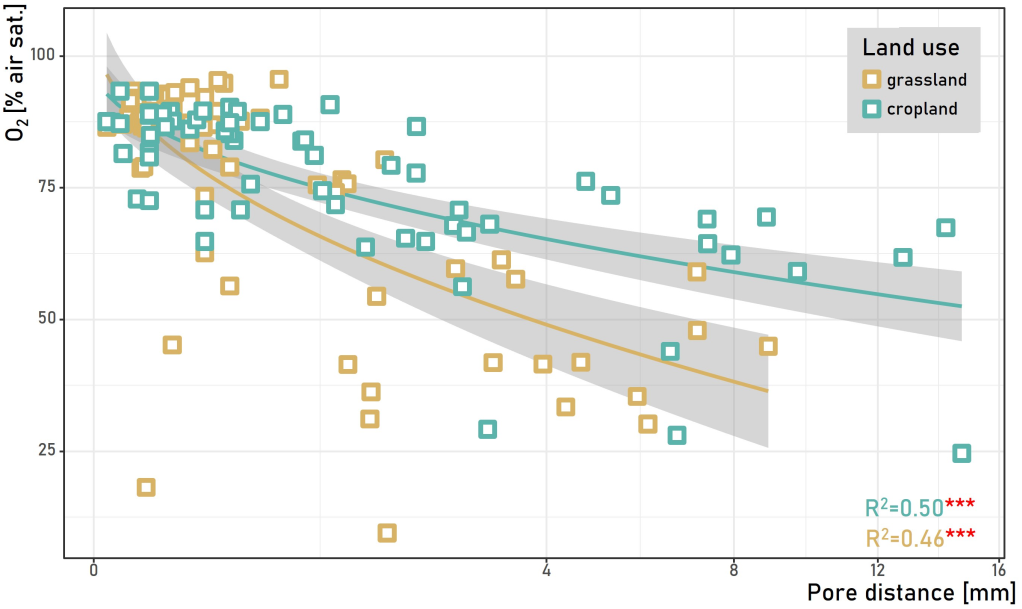


Figure S7: Association of the pore distance at the sensor tip to the next air-filled pore with the $O_2$-saturation. The solid lines represent the fitted linear models. Respective $R^2$ and significance level are given in the corner of the graph.

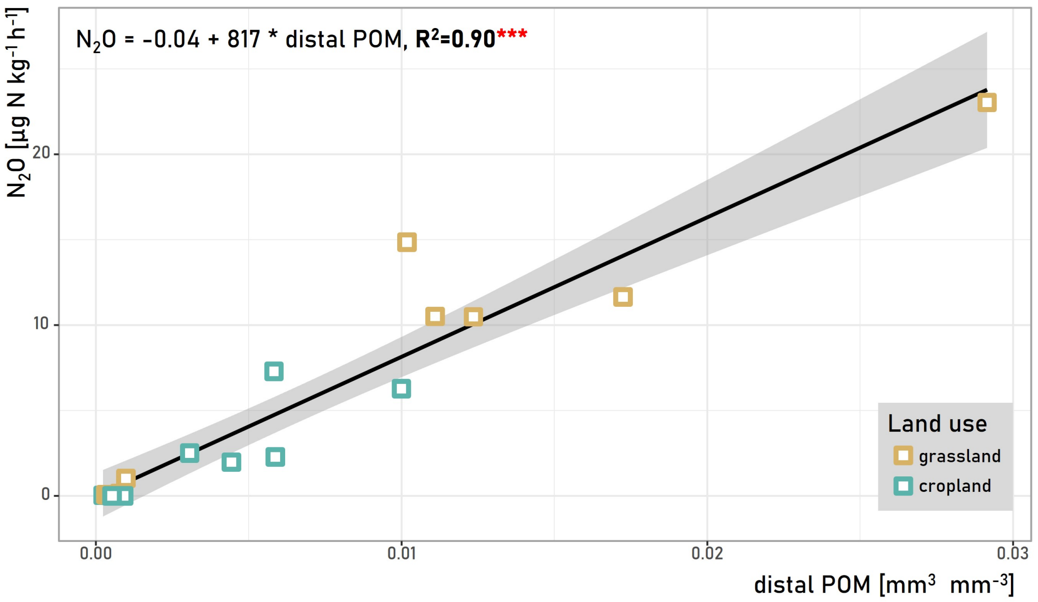


Figure S8: Association of the distal POM with $N_2O$-fluxes. Distal POM is denoted as the amount of POM >0.6 mm away from air-filled pores.*** marks a p-values <0.001.

#### 1.2 Tables

**Table S1.** Mean values and standard deviation segmented classes derived from X-ray CT scanning of 3 cm cores the two land uses under field structured conditions. Different letters indicate significant differences as derived from an LSD test comparing all groups of land use types and moisture contents (n=3, p value <0.05). Note that the image voxel size was 19*19*19 µm. Thus smaller water-filled pores and POM could not be detected.

| | Grassland | | | Cropland | | |
|---|---|---|---|---|---|---|
| matric potential [%] | 160 | 40 | 8 | 160 | 40 | 8 |
| air-filled pores [%] | 18.5±6.32a | 14.4±3.38ab | 10.3±3.73b | 13.1±2.55ab | 10.5±1.97b | 10.7±0.66b |
| water-filled pores [%] | 2.0±1.66ab | 2.2±1.34ab | 3.0±0.32a | 0.4±0.07c | 0.3±0.02c | 1.0±0.03bc |
| POM [%] | 4.4±0.17ab | 4.0±1.04ab | 4.6±1.96a | 1.1±0.43c | 1.0±0.33c | 2.7±1.26bc |
| stones [%] | 4.7±0.54a | 4.9±0.17a | 5.1±0.78a | 2.8±0.22b | 2.9±0.32b | 3.3±0.13b |
